# A Framework for Autonomous AI-Driven Drug Discovery

**DOI:** 10.1101/2024.12.17.629024

**Authors:** Douglas W. Selinger, Timothy R. Wall, Eleni Stylianou, Ehab M. Khalil, Jedidiah Gaetz, Oren Levy

## Abstract

The exponential increase in biomedical data offers unprecedented opportunities for drug discovery, yet overwhelms traditional data analysis methods, limiting the pace of new drug development. Here we introduce a framework for autonomous artificial intelligence (AI)-driven drug discovery that integrates knowledge graphs with large language models (LLMs). It is capable of planning and carrying out automated drug discovery programs at a massive scale while providing details of its research strategy, progress, and all supporting data. At the heart of this framework lies the focal graph - a novel construct that harnesses centrality algorithms to distill vast, noisy datasets into concise, transparent, data-driven hypotheses. We demonstrate that even small-scale applications of this highly scalable approach can yield novel, transparent insights relevant to multiple stages of the drug discovery process, including chemical structure-based target prediction, and present the implementation of a system which autonomously plans and executes a multi-step target discovery workflow.

**Graphical Abstract:** 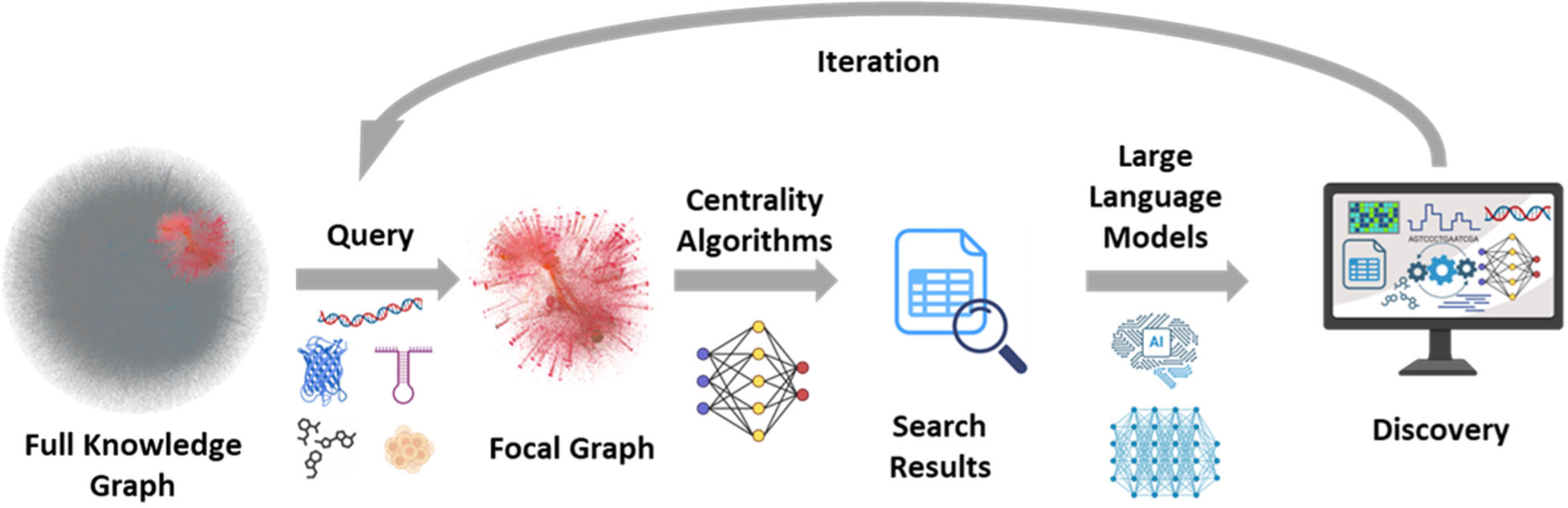

## Introduction

In recent decades, biomedical research has entered an era marked by unprecedented data generation. Advances in multi-omics, chemical biology, and high-throughput screening have led to a deluge of information, creating significant data management and interpretation challenges. This trend is exemplified by Eroom’s Law, which observes that drug discovery has become slower and more expensive over time, despite technological progress (*1*, *2*). At the same time, the vast stores of untapped biomedical data offer a potential solution to this trend (*3*), yet the sheer volume, noise, and complexity of these datasets have rendered them largely inaccessible through traditional research approaches (*4*, *5*). There is tremendous interest in the potential of AI-driven solutions to overcome these barriers, however many challenges remain (*6*, *7*).

Here we propose a framework for autonomous AI-driven drug discovery, capable of navigating and integrating large-scale, complex datasets to uncover novel scientific insights. At the core of this system is a novel subgraph generated from a specific query of a larger knowledge graph which we have named a “focal graph” (*8*). Unlike traditional knowledge graphs, which grow unwieldy as they expand, focal graphs leverage centrality algorithms, such as PageRank, to focus on highly connected subregions within larger networks (*9*). This targeted approach enables the focal graph to identify connections between key nodes - such as compounds, genes, or biological pathways - relevant to specific research queries, facilitating efficient data extraction even from vast, noisy datasets.

Building on this foundation, we will show that such a system can employ large language models (LLMs) to autonomously plan, execute, and interpret focal graph-based searches. LLMs can function as both search agents and interpreters, capable of generating hypotheses, refining search strategies, and summarizing findings across thousands of queries. We propose that this integrated approach allows continuous, iterative discovery, where new insights inform subsequent searches, and where the results can be synthesized into actionable conclusions. The autonomous capability of the system has the potential to scale effortlessly, transforming the biomedical data deluge into a torrent of structured, evidence-backed insights.

Here, we demonstrate the application of this focal graph-based, AI-driven system to multiple stages of the drug discovery process. By examining cases where focal graph analyses reveal novel insights, we show that even small-scale applications yield data-supported hypotheses. Moreover, we illustrate how scaling this approach across larger datasets amplifies its power, making it feasible to conduct research programs that would be impractical for human scientists alone. While ultimately limited and biased by available data, focal graphs can operate on a far wider set of data sources than literature-only based approaches and, because of the use of consensus, work robustly on noisy data that can present challenges to purely machine learning-based approaches. As data-driven methodologies become essential in modern drug discovery, our approach offers a promising path toward autonomous, AI-driven drug discovery—one that combines AI scalability with the transparency and rigor required for scientific discovery.

## Focal Graphs

Knowledge graphs are structured representations of real-world entities, with the entities represented as nodes, e.g., circles, and their relationships represented as edges, e.g., lines or arrows. While they have become increasingly prevalent in drug discovery and biomedical research (*10–12*), their utility is often limited by computational complexity and interpretability challenges when applied to complex diseases and drug mechanisms (*13*). As we will show, focal graphs address these limitations in multiple ways.

Focal graphs are subgraphs of knowledge graphs, constructed such that the properties of their seed members, or “queries”, can be inferred from their graph properties. Many kinds of focal graphs can be constructed to answer many different kinds of questions. For example, focal graphs constructed based on the known targets of structurally related compounds can be used to infer the putative target(s) of a seed compound (*8*, *14*). They can be adapted to many other applications by modifying their construction and analysis, for example by varying the seed type, the knowledge graph from which they are derived, or by optimizing them to infer a particular property or set of properties. This makes them an extremely flexible tool for extracting many different kinds of insights.

Starting from a query of interest - whether a compound, gene list, or disease signature - a focused subgraph is constructed that captures the most relevant relationships within the larger knowledge network (Figure 1A). This focused extraction enables detailed analysis of specific research questions while maintaining computational tractability, even as the underlying knowledge graph grows arbitrarily large. Centrality algorithms are mathematical methods used to determine the relative importance or influence of nodes within a graph based on their connectivity and position. Their application to focal graphs yields concise, rank-ordered lists of the most highly connected graph members, which can be used in turn to infer the potential properties of the original queries.

**Figure 1.**
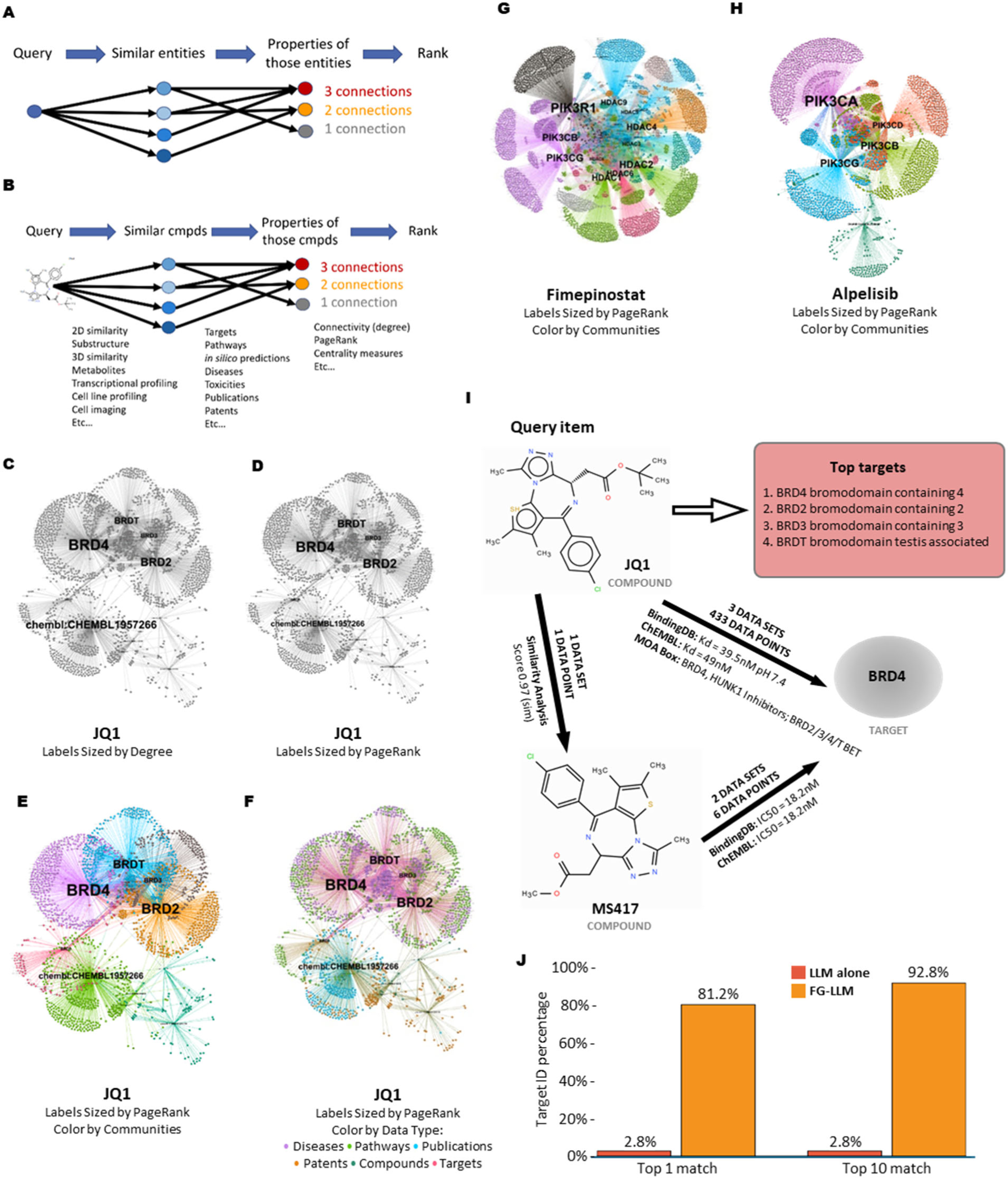
Overview of focal graphs. A) General focal graph-building workflow showing a query node (leftmost blue circle) linked by edges (represented by arrows) to nodes representing similar entities (middle circles in shades of blue), then linked to nodes representing their properties (rightmost circles), which are then ranked by a graph centrality algorithm, such as degree of connectivity. B) Focal graph building workflow for compound target ID. The query example here, JQ1, is a compound structure from which similar compounds can be found and additional information on its targets, pathways, disease indications and other properties can be inferred. C,D) A focal graph generated for JQ1, a BET family inhibitor with known targets BRD4, BRD3, BRD2, & BRDT, with labels sized by degree (C) or by PageRank (D). E,F) Focal graphs for JQ1 with labels sized by PageRank and colored by graph community (E) or by data type (F). G) Focal graph of Fimepinostat, a pan-PI3K and pan-HDAC inhibitor, with labels sized by PageRank and colored by graph community. H) Focal graph of Alpelosib, an alpha specific PI3K inhibitor, with labels sized by PageRank and colored by graph community. Focal graph images in C-H were generated by Gephi (https://gephi.org). I) Selected focal graph evidence https: J) Percentage of compounds (out of 500) for which the top target predictions of the FG-LLM (orange) or LLM alone (red) contain at least one of the annotated targets. See Supplementary Methods for details.

The construction of a focal graph follows a systematic workflow (Figure 1A). First, entities similar to the query are identified, creating an initial layer of relationships. These similar entities are then connected to their associated properties - such as compounds, targets, pathways, or disease associations - forming a second layer of connectivity. The resulting multi-layered network can then be analyzed using centrality algorithms, such as PageRank, which have proven effective in identifying essential nodes in biological networks (See Supplementary Methods)(*15*).

Centrality algorithms identify results with the broadest levels of support, making them highly robust to noise. They also focus attention on the most strongly supported results, making them highly concise and scalable. The most experimentally supported results are ranked first, regardless of how much data is searched or how many additional results are returned. In fact, the larger and more diverse the global knowledge graph becomes, the better the focal graph-derived results should be, without sacrificing robustness, concision, transparency, or simplicity. Any structured data source or database could be included as part of a knowledge graph from which focal graphs are derived, including large, diverse resources such as Open Targets (*16*), Gene Expression Omnibus (GEO) (*17*), and ChEMBL (*18*, *19*), allowing for connections within and between them, potentially revealing deeper insights.

Focal graphs are exceptionally robust to noise. Datasets with high levels of random noise will fail to agree with other data sources, pushing their results lower in the rankings. Datasets with systemic noise will only agree with other data with the same bias, but not with other, independent approaches, pushing their results lower in the rankings while also potentially making the source of the systematic noise easier to identify.

In contrast to many machine learning (ML) based methods, the data supporting focal graph-derived results can be easily identified and traced back to their original sources, allowing in-depth evaluation by subject matter experts. This high level of transparency is critical for scientific and regulatory applications where specific details of the supporting data is crucial for decision-making. This combination of properties makes focal graphs ideally suited for dealing with the huge explosion of complex, large scale datasets that are ubiquitous in modern biomedical research.

The power of focal graphs becomes evident when examining specific examples. A focal graph generated using centrality algorithms for the BET bromodomain inhibitor JQ1 (Figure 1B), a well-characterized chemical probe (*20*), reveals distinct communities of interconnected nodes sized by degree (Figure 1C), by PageRank (1D), or colored by graph community (1E) or data type (1F), each representing different aspects of the compound’s biological activity. These graph communities, or neighborhoods, share similar connections and potentially also biological functions, and can be excised and studied independently. Community structure can be driven by many factors including common chemical structure, biological processes, data provenance, bioactivity, or some combination of these.

When colored by data type (Diseases, Pathways, Publications, Patents, Compounds and Targets, Figure 1F), we can observe how different types of evidence contribute to an understanding of the compound’s mechanism of action. Generating a focal graph using the structure of JQ1 as a query identifies connections to its known molecular targets, including BRD4, BRD2, BRD3 and BRDT (Figure 1I). This illustrates how focal graphs can integrate both direct and indirect supporting data, e.g., chemical similarity algorithms and data from diverse sources such as ChEMBL and PubChem (*18*, *19*, *21*). Focal graphs constructed for Fimepinostat, a pan-PI3K and pan-HDAC inhibitor (Figure 1G) (*22*), and Alpelisib, an alpha specific PI3K inhibitor (Figure 1H) (*23*), reflect each compound’s known, distinct polypharmacology.

## Synergy with Machine Learning Approaches

While machine learning approaches have demonstrated impressive predictive capabilities in drug discovery (*24*), they often face challenges with data heterogeneity, lack of transparency, result interpretability, and the need for large training sets (*25*). Neural networks in particular, including graph neural networks and deep learning models, typically operate as “black boxes,” making it difficult to trace the evidence supporting their predictions (*26*). In contrast, focal graphs preserve the provenance of their insights, allowing researchers to examine the exact experimental data and relationships that support each conclusion.

Many of the edges of the focal graphs in Figures 1-3 were created at run time by chemical similarity algorithms. Edges can in principle be generated by any kind of algorithm, including ML-based predictions (*27*, *28*), allowing focal graphs to simultaneously integrate diverse data sets and *also* a variety of algorithms, identifying for example, where experimental data and predictive algorithms most converge. This can help optimize algorithms by identifying where they agree, or fail to agree, either with other algorithms or with experimental data. Agreement with experimental data may provide a broader view of the biological or chemical context that drove the algorithmic prediction, making it more interpretable and potentially yielding insights that may help refine the predictive approach.

The robustness of focal graphs stems from their ability to identify consensus across multiple data sources and experimental approaches. Unlike supervised learning methods that may learn spurious correlations from training data (*29*), focal graph centrality algorithms naturally highlight instances where multiple independent lines of evidence converge on the same conclusion. This approach is particularly valuable in drug discovery, where experimental noise and systematic biases are common challenges (*30*).

Focal graphs also offer distinct advantages in handling sparse or novel data. While machine learning models struggle with predictions outside their training distribution, focal graphs can surface meaningful relationships for novel entities by connecting them to a small number of biologically or chemically related examples. In fact, even a single strong connection can be enough for a focal graph to generate a compelling hypothesis, whereas a ML-based model typically takes a large number of examples to train and test. This capability is especially relevant for emerging therapeutic modalities or newly discovered disease mechanisms where training data may be limited.

The relationship between focal graphs and machine learning approaches is more complementary than competitive. Machine learning can be incorporated into focal graph construction strategies by the inclusion of computationally created edges. Focal graphs can provide interpretable context for machine learning predictions by connecting them to relevant experimental data as well as predictions from other computational approaches (*31*).

Furthermore, focal graphs can themselves be used to generate training sets for machine learning models, for example by providing rank-ordered consensus examples for a variety of query types, such as a set of compound structures most associated with inhibition of a particular target or a set of genes most associated with a toxicity. Thus, the integration of focal graphs with ML-based prediction, particularly when combined with large language models, can lead to natural synergies that produce novel findings while maintaining scientific rigor and transparency.

## Focal Graph - Large Language Model (FG-LLM) Integration

Large language models (LLMs) such as ChatGPT or Claude have a remarkable ability to converse in natural language, incorporating an impressive breadth of information gleaned from massive text-based training sets. Expanding their capabilities beyond text-based data is an area of active investigation (*32*, *33*). Combining LLMs with an internet search engine has proven to be a highly successful and easily implemented strategy (*34*). However, internet searches do not currently include the experimental results from large scale biomedical datasets, such as chemical structure/bioactivity data or multi-omics datasets, and it remains unclear the extent to which LLMs can interpret chemical structures in a way that enables chemical similarity or substructure searching (*35*, *36*). A focal graph search is an effective way to connect these data to LLMs, as demonstrated in the use cases described below, potentially reducing the frequency of hallucinations and dramatically enhancing the power of LLMs to advance drug discovery.

Integration of focal graphs with large language models allows the automatic generation, execution, and interpretation of searches, allowing them to be included as part of large scale “intelligent” workflows. These automated searches can be run serially for many iterations, potentially allowing execution of an extended computational research program that uses massive amounts of existing data - all while maintaining full transparency with respect to the research strategy and the data that supports the hypotheses uncovered. Key findings of large scale workflow runs can be summarized for easy evaluation and provide the basis for larger scale target prioritization efforts. We will return to this topic and cover it in more detail later.

We designed a quantitative benchmark for chemical structure-based target identification to evaluate the FG-LLM approach (see Supplementary Methods). To increase its relevance for real-world applications where the targets of compounds are unknown, we created a modified reference set, based on the MoA Box (*37*), where each compound was modified by the addition of a single fluorine atom. This change is small enough that the compounds’ targets are unlikely to be changed, however it forces the system to rely on chemical structure-based inferences rather than on trivial target lookups, which would not be possible in any case when analyzing novel compounds.

We applied an FG-LLM approach to a set of 500 benchmarking compounds, deriving a focal graph for each, and prompting the LLM to use each set of focal graph results to infer the most likely target. In parallel, we used a highly similar prompt to direct the same LLM to perform the same task, but without providing focal graph results. Comparing the inferred targets from these two approaches with the benchmark targets, we found that the single top target inferred by the FG-LLM approach matched a benchmark target in 81.2% (406/500) of cases, and the top 10 inferred targets by FG-LLM contained a match in 92.8% (464/500) of cases. In contrast, the LLM without focal graph results matched a benchmark target for only 2.8% (14/500) of compounds whether the top 1 or top 10 results were considered.

An analysis of discordant FG-LLM results (see Supplementary Methods and Supplementary Table S6) suggests ways the system could be further improved, such as by the inclusion of additional chemical fingerprints, and points to the potential utility of focal graphs for identifying errors and omissions in manually curated annotations, which could lead to improvements in completeness, uniformity and reliability. The performance of FG-LLM approaches across other chemical and biological benchmarks, potentially in combination with additional specialized scientific tools, represents a promising direction for future research.

## Diverse Applications of Focal Graphs to Drug Discovery

Focal graphs can capture complex relationships between biological entities or nodes, making them applicable to the exceptionally wide range of datasets characteristic of modern biomedical research. This allows them to find connections between almost any data type and makes them applicable to a wide range of experimental approaches, algorithms, and scientific questions (*8*, *14*, *38*).

Here we present several case studies that demonstrate the versatility and power of focal graphs. We show how large language models can be employed to systematically interpret their results and suggest new avenues for investigation. The examples presented here, while focused on specific use cases, illustrate general principles that can be applied to a broad range of drug discovery challenges.

### Chemistry-Driven Target Discovery

Phenotypic screening is widely used as an approach to discover highly relevant modulators of biological processes for drug development. However, identifying the molecular targets of the screening hits and the mechanisms involved, remains a difficult challenge (*39–42*).

We used a focal graph to analyze 23 compounds from a chemical series (series 46) identified by Calderon et al. (*43*) as having antimalarial activity but whose target(s) are unknown (Figure 2). A focal graph search of the compound series revealed two highly ranked potential targets: poly(ADP-ribose) polymerase 1 (PARP1) and dihydroorotate dehydrogenase (DHODH), each supported by multiple lines of evidence (Figure 2A-D). While the focal graph search identified more high potency series 46-analogs connected to PARP1 than to DHODH, the LLM analysis found that DHODH was more strongly connected to malaria, with data coming from multiple sources. These included the connection to the DHODH inhibitor DSM265, an investigational antimalarial treatment that progressed to phase II clinical trials (*44*, *45*). This broader, cross-data type perspective led the LLM to conclude that the antimalarial activity of the series was best explained by DHODH inhibition (Figure 2E). Notably, the focal graph identified a publication that described a structurally related series as having dual PARP1 and DHODH activity (*46*), suggesting that series 46 may in fact inhibit both targets.

**Figure 2.**
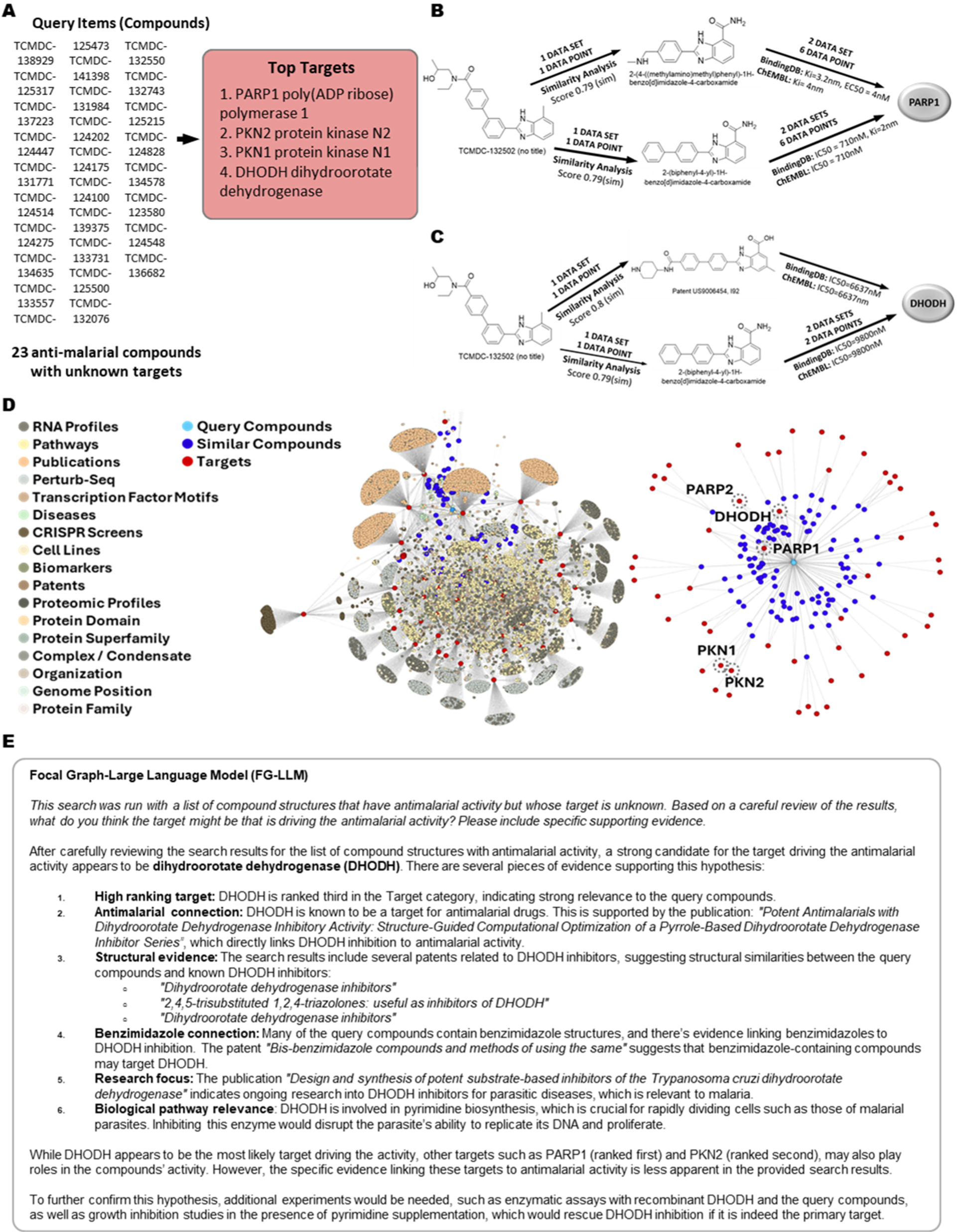
Focal graph analysis of a 23-member compound series, active in an antimalarial phenotypic screen. A) The top ranked targets from a focal graph search of 23 compounds from an antimalarial phenotypic screen are shown. B,C) Selected focal graph evidence paths connecting a member of the compound series to PARP1 (B) or to DHODH (C) through chemical similarity and binding data. D) The focal graph of the 23 antimalarial compounds showing connections to many diverse data types. A subgraph of the focal graph is shown on the right comprising the connections between only the query compounds, similar compounds, and targets. The top ranked targets are labeled. E) Large Language Model (LLM) analysis of the focal graph search which highlights DHODH inhibition as the best-supported antimalarial target hypothesis (48, 49). Focal graph images in D were generated by Gephi (https://gephi.org). Focal graphs were used in combination with Claude 3.5 Sonnet, a foundation LLM from Anthropic.

This example illustrates several key advantages of the focal graph approach. The strength of the structural similarity across the full series can be analyzed in detail and its consistency (or lack thereof) with previously established structure activity relationships can be assessed. Also, because the focal graph integrates extremely diverse data types, many aspects of the target hypothesis can be evaluated at once. This includes considering not only the molecular pharmacology but also the biology of the target, as described in the scientific literature and/or inferred directly from large scale data sets present in the focal graph. Ultimately, the transparency of the results allows expert evaluation of the full breadth of the underlying data and enables a nuanced analysis tailored to the specific goals and detailed context of the overall project.

A focal graph approach was used recently to predict DPP4 and HSD17β13 as direct molecular targets of α-Terthienyl, a compound identified in a phenotypic screen and found to significantly reduce hepatic steatosis in a murine model. Both targets were subsequently confirmed experimentally in biochemical assays, highlighting the ability of focal graphs to predict the molecular targets of novel, therapeutically relevant compounds (*47*).

### Morphological Profiling

Cell painting, a high-dimensional imaging assay that captures cellular morphological states, offers an unbiased approach to characterization of a compound’s mechanisms of action (*50–52*). However, translating these rich phenotypic profiles into specific molecular hypotheses remains challenging, though significant progress with this methodology is being made (*53*). We selected a cluster of 10 compounds with cell painting profiles most similar to the HDAC inhibitor vorinostat by UMAP analysis using the Ardigen phenAID JUMP-CP Data explorer (Figure 3A)(*54*). Using the structures of the compounds in this cluster as a focal graph query revealed histone deacetylases (HDACs 1,6,3 and 2) as the top-ranked targets (Figure 3B). This is based not only on the known targets of vorinostat, but also on the activities of other compounds in the cluster and their structurally related analogs (Figure 3C,D).

**Figure 3.**
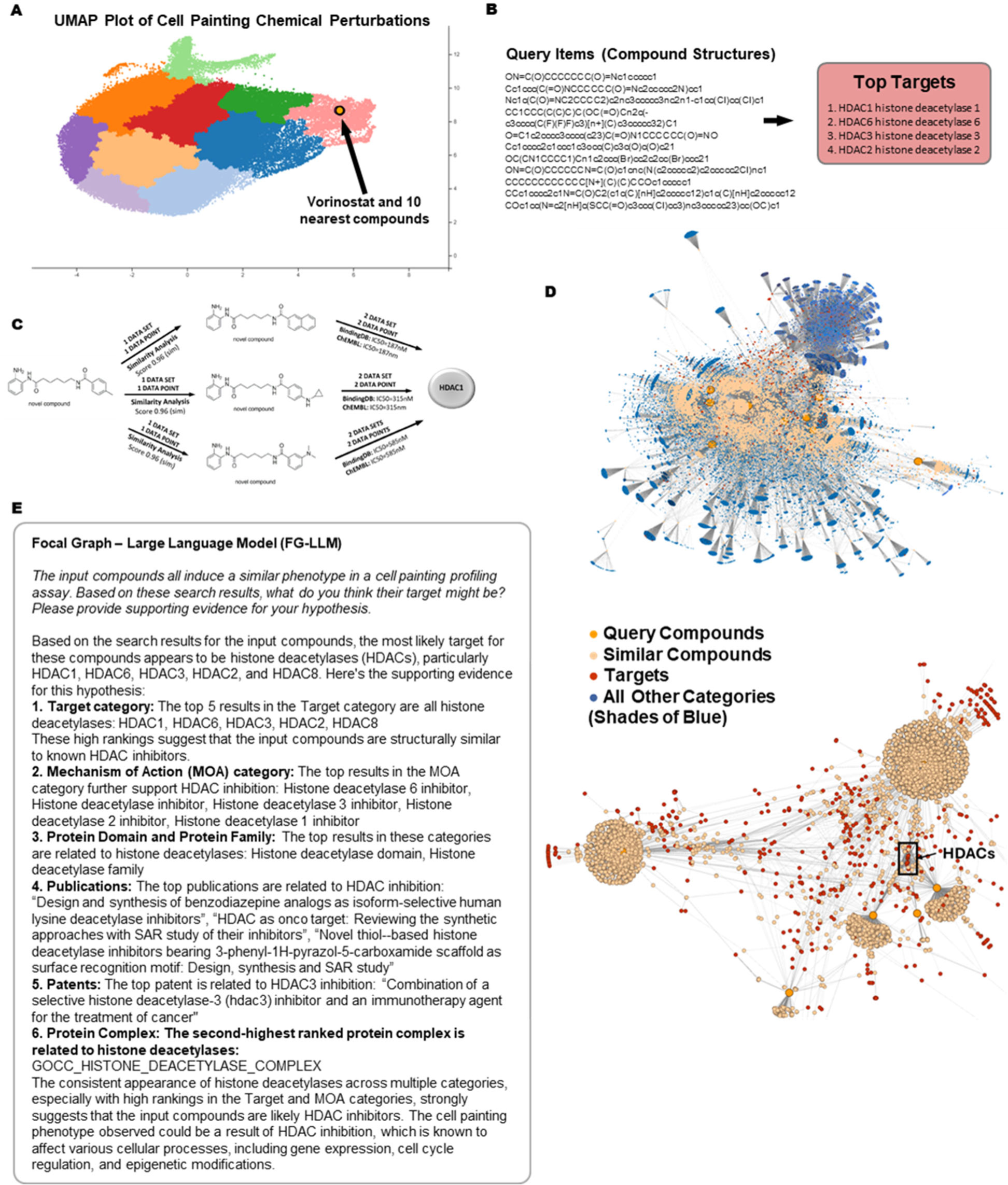
Focal graph analysis of a cell painting compound cluster. A) Selection of a cluster of 10 compounds with cell painting profiles most similar to Vorinostat by UMAP analysis using the Ardigen phenAID JUMP-CP Data Explorer (https://phenaid.ardigen.com/jumpcpexplorer/). The x and y axes represent the first two dimensions of the UMAP projection, respectively. B) Top ranked targets from a focal graph search with the Vorinostat cell painting cluster, including Vorinostat itself. SMILES of the compounds in the cluster are shown. C) Selected focal graph evidence paths connecting a member of the compound series to HDAC1. D) The full vorinostat focal graph (top) and a subgraph (below) showing only query compounds, similar compounds, and targets. The location of a cluster of top ranked HDACs are highlighted. E) Large Language Model (LLM) analysis of the focal graph search which highlights HDAC inhibition as the best-supported target hypothesis for this compound cluster (55–57). Focal graphs were used in combination with Claude 3.5 Sonnet, a foundational LLM from Anthropic.

The focal graph visualization (Figure 3D) illustrates how relationships between the query compounds and HDAC family members were identified. Multiple independent lines of evidence connect the cluster of 10 compounds to HDAC1 (Figure 3C,D), including direct binding evidence and structural similarities to known HDAC inhibitors. This convergence of evidence from diverse sources provides strong support for HDAC inhibition as the shared mechanism of this set of compounds. This was summarized by a large language model analysis of the focal graph results (Figure 3E).

We also recently used focal graphs to analyze sets of genetic perturbations that were found to cluster together in a large cell painting dataset from the JUMP-Cell Painting Consortium, uncovering both expected and unexpected relationships between genes (*38*). These results helped validate the dataset and uncovered new connections, identifying potential new drug targets that may play important roles in cancer, mitochondrial function, and nervous system development.

### Multi-Omics Target Discovery

Gene expression profiling can provide an unbiased view of disease mechanisms, but extracting actionable insights from these complex datasets remains a challenge. We analyzed a transcriptional signature derived from psoriatic lesions compared to non-lesioned skin (Figure 4A) (*58*). The focal graph was generated using a list of differentially expressed genes, with no explicit reference to psoriasis. Nevertheless, the top-ranked results included multiple independent psoriasis gene expression studies (Figure 4A) providing strong validation of the signature’s disease specificity. Large language model analysis of the search results (Figure 4B) systematically evaluated the evidence linking the signature to various skin conditions, confirming psoriasis as the best-supported diagnosis based purely on transcriptional changes.

**Figure 4.**
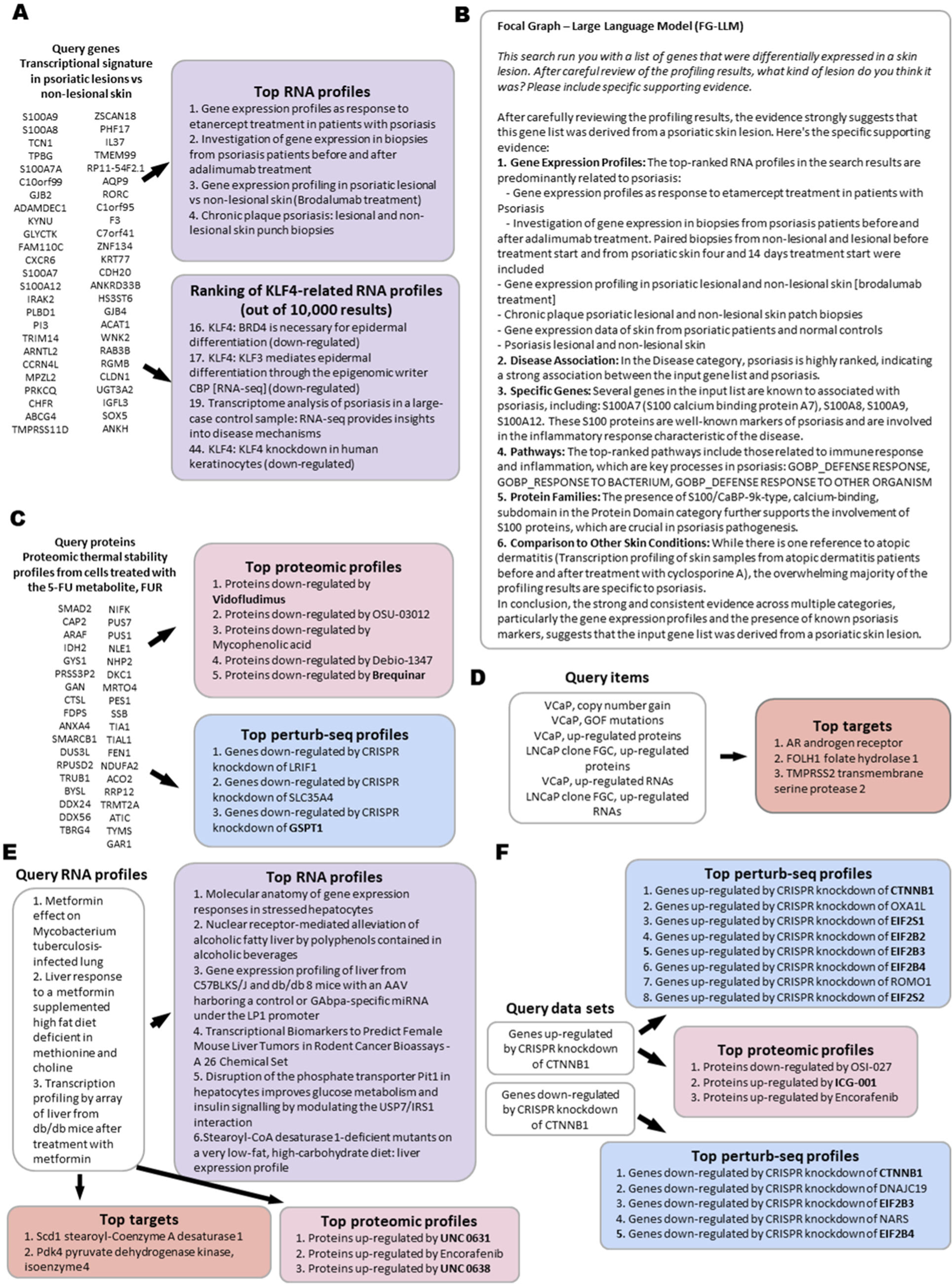
Focal graph analysis of omics profiles, discovery of novel pathway components & identification of biomarkers for precision medicine. A) RNA profile results of a focal graph search using a transcriptional signature in psoriatic lesions versus non-lesional skin as a query. The top ranked RNA Profile results are independent gene expression studies of psoriasis. Profiles of genes after modulation by KLF4 are shown. These were ranked highly (rankings shown), including 4 results in the top 50 out of 9,889 total results. B) Large Language Model (LLM) analysis of the focal graph search which highlights psoriasis as the skin lesion type most strongly supported by the data, based solely on the differential gene expression profile (58–69). C) Focal graph analysis of a CETSA proteomic thermal stability profile generated from cells treated with FUR, a 5-FU metabolite. The top proteomic profiles identified by the focal graph are shown. Two of the matching proteomic profiles are ones following treatment with DHODH inhibitors vidofludimus and brequinar. The top perturb seq profiles revealed by the focal graph included genes downregulated by CRISPR knockdown of GSPT1. D) Identification of biomarkers associated with prostate cancer using the molecular profiles of two prostate cancer cell lines VCaP and LNCaP as a query. The top targets identified were three of the most important genes linked to prostate cancer. E) 3 RNA profiles from mice treated with metformin were used as a query for a focal graph search. The top targets are strongly linked to glucose and fatty acid homeostasis. The top 100 genes from the RNA profile focal graph search were mapped to their human and rat orthologs and a signature representing all 3 species was run as another focal graph search. The results show links to RNA profiles related to glucose and fatty acid homeostasis. The top proteomic profiles show links to proteins up-regulated by small molecules UNC 0631 and UNC 0638, which are both inhibitors of EHMT2, which has been shown to be down-regulated by metformin. F) Discovery of novel components in the β-catenin pathway by focal graph analyses of genes up- or down-regulated after CRISPR knockdown of β-catenin. In each case the top Perturb seq profile results included multiple subunits of eukaryotic initiation factor 2 complex (eIF2). The top proteomic profiles contain a profile of proteins up-regulated by ICG-001, a known small molecule β-catenin inhibitor. Focal graphs were used in combination with Claude 3.5 Sonnet, a foundational LLM from Anthropic.

Further analysis of the focal graph results revealed an intriguing connection to KLF4, with multiple KLF4-modulating profiles showing significant overlap with the disease signature (Figure 4A). KLF4, a transcription factor known to regulate epidermal differentiation (*59*), appeared in four of the top 50 results out of nearly 10,000 total profiles (the ranking of each KLF4 RNA profile in the top 50 results is shown). This enrichment suggests that KLF4 modulation could potentially reverse the disease signature, making it a promising therapeutic target. The hypothesis is particularly compelling given KLF4’s established role in maintaining skin barrier function and regulating inflammatory responses (*59*, *60*).

This example illustrates how focal graphs can generate novel therapeutic hypotheses by integrating disease signatures with large-scale perturbational datasets. The approach not only validated the disease specificity of the signature but also identified a potential therapeutic target that might have been overlooked by traditional analysis methods. The transparent nature of the focal graph results allows direct evaluation of the evidence supporting the KLF4 hypothesis, from its effects on specific psoriasis-associated genes to its broader role in skin biology.

### Proteomic Profiling

Focal graphs can be used to analyze complex proteomic datasets. This is illustrated by analysis of proteomic thermal stability profiles generated from cells treated with the 5-FU metabolites FUR and FUDR and subjected to a cellular thermal shift assay, CETSA (Figure 4C) (*69*). 5-FU, a cornerstone of cancer chemotherapy for over 50 years, has long been known to inhibit thymidylate synthetase, but questions remain about its mechanism of action and strategies to improve its anti-cancer efficacy (*70*, *71*). A focal graph analysis confirmed 5-FU’s known effects on pyrimidine synthesis, and in addition, identified connections to proteomic profiles matching those following treatment with two DHODH inhibitors vidofludimus and brequinar (Figure 4C).

DHODH inhibition has been reported to be synergistic with 5-FU treatment in cancer models (*72–74*). DHODH is an enzyme in the pyrimidine biosynthesis pathway which produces uracil, providing a clear mechanism by which DHODH inhibitors may be synergistic with 5-FU treatment.

Intriguingly, the analysis revealed unexpected similarities between FUR/FUDR-induced proteomic changes and those caused by CRISPR knockdown of GSPT1 (Perturb-seq profile in Figure 4C), suggesting interference with protein translation termination. This connection is particularly noteworthy due to the recent emergence of GSPT1 as a novel clinical stage oncology target for which PROTACs have been developed (*75*).

The CETSA example demonstrates the ability of the focal graph approach to weave together diverse data types, in this case connecting compound-proteomic interaction data (CETSA) to compound-induced proteomic protein level changes (MoA Atlas data)(*76*) and to transcriptomic RNA level changes after CRISPR knockdown (Perturb-Seq data)(*77*).

In summary, a focal graph analysis revealed additional data supporting possible synergy between 5-FU and DHODH inhibitors. It also identified a connection to GSPT1 knockdown, suggesting that the disruption of translational termination may be a part of 5-FU’s mechanism and pointing to potential synergy between 5-FU and GSPT1 targeted therapeutics.

### Precision Medicine

The development of robust patient stratification strategies such as the identification of biomarker signatures, remains a critical focus in precision oncology. Here we demonstrate how focal graphs can accelerate biomarker discovery by uncovering the molecular mechanisms that underlie a therapeutic response. A focal graph was queried with the molecular profiles of two prostate cancer cell lines (VCaP and LNCaP) that include RNA expression, protein levels, copy number alterations, and mutations (*78*, *79*). The focal graph identified the most biologically significant genes associated with these cell lines as AR (androgen receptor), FOLH1 (PSMA), and TMPRSS2 (Figure 4D). These genes are arguably the most important, extensively validated and clinically relevant genes associated with prostate cancer. AR status guides treatment selection, FOLH1 expression enables PSMA-targeted imaging and therapy, and TMPRSS2 fusions serve as diagnostic and prognostic markers (*80–82*).

This example illustrates a generalizable strategy for mechanism-based biomarker development. Lists of cell lines showing differential response to a therapeutic agent, can be analyzed by focal graphs to identify the molecular features that distinguish responders from non-responders. By analyzing large-scale cell line characterization datasets like CCLE and DepMap (*78*, *83*, *84*), this approach can reveal not just correlative biomarkers but the underlying biological drivers of therapeutic sensitivity. The resulting mechanistic signatures can then be used to identify additional cell lines with similar molecular profiles, enabling rapid experimental validation of both the biomarker and its hypothesized mechanism of action.

The progression from response pattern to mechanistic understanding is particularly powerful for novel therapeutics where mechanisms are usually not immediately apparent (*85*). The multi-omic nature of the analysis helps capture the full biological context of the drug response, while the transparent nature of focal graph analysis ensures that the proposed mechanisms are supported by clear experimental evidence. This mechanism-first approach could significantly accelerate the development of companion diagnostics by providing patient selection strategies with a solid mechanistic basis.

### Safety Assessment

Focal graph analyses that are based on many types of queries, including chemical structures, multi-omics data, and cell line profiles, can be used to generate mechanism-of-toxicity hypotheses for safety assessment. Focal graphs starting from chemical structure were used to investigate the mechanism of drug-induced skin toxicity in cynomolgus monkeys (*14*). The test compound was found to be structurally similar to compounds with a similar cell line proliferation profile to PI3K inhibitors. Many PI3K inhibitors are known to cause skin toxicity, suggesting a possible role for the PI3K pathway in the toxicity of the tested compound (*86*).

Omics profiling, such as the psoriasis-derived RNA signature described previously, can be used to match a drug response to one or more profiles of a perturbagen with a described mechanism, which can then be used to identify potential mechanism(s) underlying the toxicity. A list of cell lines sensitive or resistant to a compound, such as those from a PRISM analysis (*87*), can be used as a focal graph query to identify drug targets or mechanisms that may be driving toxicity. Many independent focal graph searches can be run in parallel, with mechanistic hypotheses assessed based on the strength of their supporting data from individual analyses, as well as agreement across analyses. This enables a multi-faceted, data-driven, comprehensive approach to safety assessment.

### Biomarker Development

The development of pharmacodynamic biomarkers from gene expression signatures is another application of focal graphs. A focal graph analysis identified 100 mouse genes that were most commonly modulated across 3 independent transcriptional profiles of mice treated with metformin, a widely prescribed drug for the treatment of diabetes (*88–90*). Notably, the top 2 genes in the signature, Scd1 and Pdk4 (Figure 4E), are known to be involved in glucose and fatty acid homeostasis. The 100 gene mouse signature, supplemented with the corresponding human and rat orthologs, was used in a subsequent focal graph search (Figure 4E), yielding top connections to other, independent transcriptional profiling studies related to the impacts of diet on hepatic function, including effects on insulin signaling and fatty liver. This suggests that this signature indeed captures the expected biology and that this strategy could be used as part of an effort to develop a pharmacodynamic biomarker for metformin response. Interestingly, two of the top three proteomic profiles overlapped the input signature by 6 of 100 genes. These are the profiles of UNC 0631 & UNC 0638, which both inhibit EHMT2, a histone lysine methyltransferase (*91*, *92*). Metformin has been reported to down-regulate EHMT1 & EHMT2 expression at the RNA and protein levels (*93*).

This example illustrates how focal graphs can be used to rapidly develop biomarkers using gene signatures as input. In this case the signatures were derived from drug-induced RNA profiles, but in principle they could come from many possible origins, including pathways, diseases, toxicities, and genetic perturbations, or from many possible data types, including transcriptomics, proteomics, genomics, metabolomics, or a mixture thereof.

### Disease Indication Discovery

The discovery of novel disease indications for a target or a therapeutic is a further application of focal graphs in which disease processes based on gene signatures can be identified and characterized. When presented with a list of 50 up-regulated genes from a spatial, single-cell transcriptomics study of dystrophic muscle (*94*), without any explicit mention of Duchenne Muscular Dystrophy (DMD), the integrated focal graph-LLM correctly identified DMD as the disease most strongly connected to the signature (Figure 5, Supplementary Fig. S1). The system identified multiple independent RNA profiles specifically mentioning “mdx mice” - a well-established DMD model - and found studies directly comparing dystrophin-deficient and wild-type muscle tissue.

**Figure 5.**
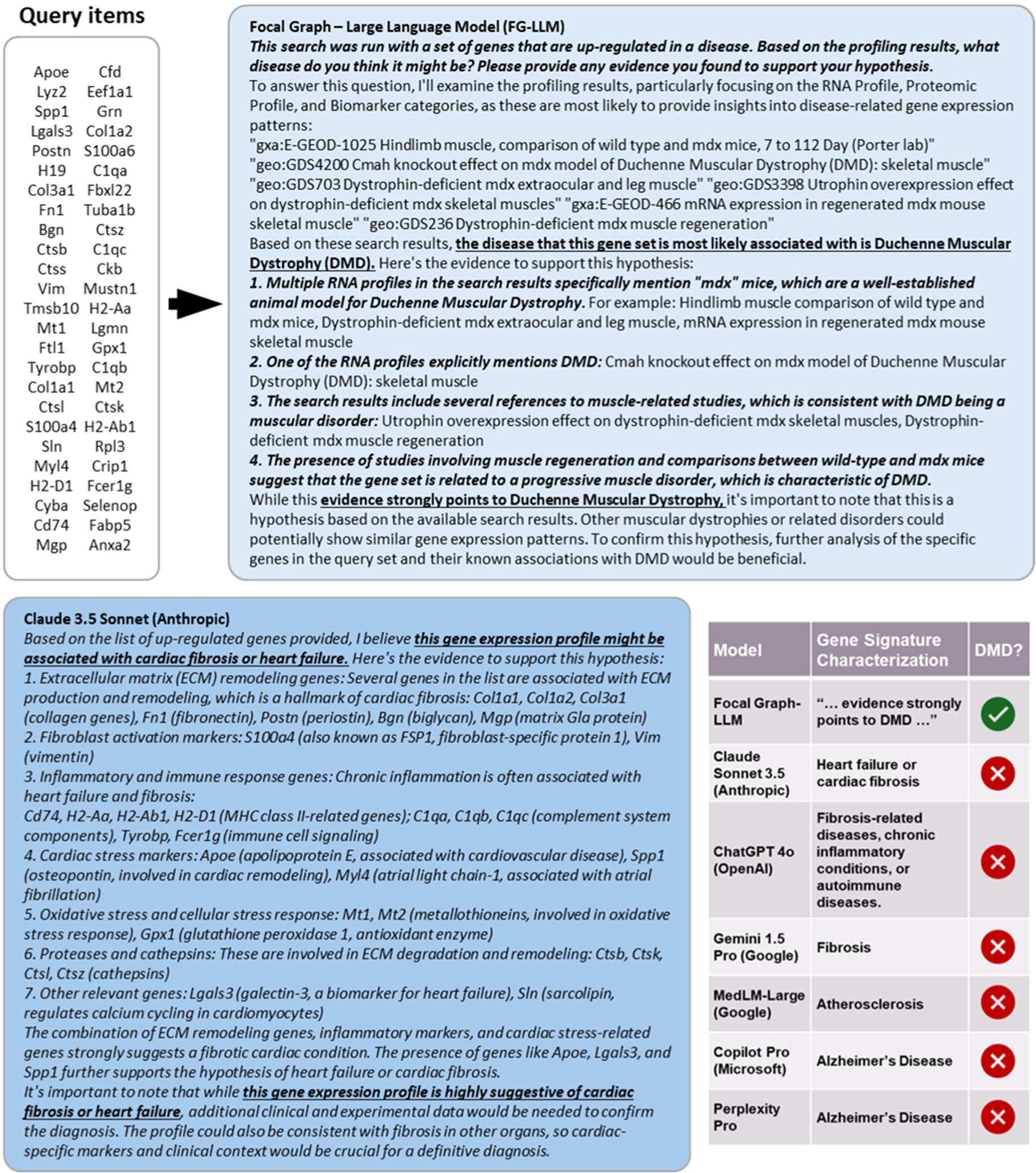
Identification of Duchenne Muscular Dystrophy from a gene signature by an LLM integrated with the focal graph. Focal graph and LLM analysis of 50 genes upregulated in Duchenne Muscular Dystrophy (DMD) without any explicit mention of DMD. The LLM integrated with the focal graph correctly identified the disease based on the focal graph search results (99–103). The same gene list was analyzed by leading LLM models alone, including Claude 3.5 Sonnet (Anthropic), ChatGPT-4o (OpenAI), MedLM-Large (Google), Gemini 1.5 Pro (Google), Co-Pilot Pro (Microsoft), and Perplexity Pro (Perplexity). None correctly identified DMD as the source of the signature attributing the gene expression pattern to cardiac fibrosis, atherosclerosis, Alzheimer’s disease, or general inflammatory conditions. Focal graphs were used in combination with Claude 3.5 Sonnet, a foundational LLM from Anthropic. Underlined emphases added. See Supplementary Fig. S1 for unabridged transcripts of the LLM responses.

This example shows how focal graphs can be used to characterize disease processes in a relatively unbiased way, enabling connections to other data that can yield insights into the underlying disease mechanism and suggest new therapeutic areas.

### Focal Graph-Integrated LLMs Outperform Non-Integrated LLMs

When the same gene list was analyzed by leading LLM models, including Claude 3.5, GPT-4, MedLM-Large, Co-Pilot Pro and Perplexity Pro, (Figure 5, Supplementary Fig. S1), none of which include a focal graph analysis, none correctly identified DMD as the source of the signature (*34*, *95–98*). Instead, these models variously attributed the gene expression pattern to cardiac fibrosis, atherosclerosis, Alzheimer’s disease, or general inflammatory conditions. This discrepancy highlights a key advantage of focal graphs: the ability to ground LLMs in specific experimental data rather than relying solely on pattern matching from training data.

The DMD example demonstrates how focal graphs can enhance LLM capabilities by providing access to specialized datasets that may not be well-represented in general training data or which may not be well suited for current LLM training methods. The use of focal graphs as a connector enables the LLM’s to synthesize and communicate complex biological insights while ensuring conclusions are supported by explicit experimental evidence. This combination of computational reasoning with empirical data validation represents a powerful new paradigm for biological data analysis. As the power and breadth of LLMs inevitably increases, so too should the power of combining them with other tools.

### Systematic Discovery of Novel Pathway Components

The prediction of novel targets that could modulate disease relevant pathways is central to the development of new therapeutic strategies. The power of focal graphs to uncover novel pathway connections is demonstrated by focal graph analysis of Wnt pathway modulation. A focal graph analysis was run using mRNAs up- or down-regulated after CRISPR knockdown of β-catenin (*77*), a central mediator of Wnt signaling, revealing unexpected connections to the translational machinery (Figure 4F). Multiple subunits of eukaryotic initiation factor 2 (EIF2) showed transcriptional responses that were highly correlated to β-catenin knockdown (Perturb seq datasets). This was found for multiple subunits of EIF2 and for both up- and down-regulated genes, suggesting that this is a robust connection.

The biological relevance of these findings was further reinforced by cross-modal validation, with strong correlations seen between β-catenin knockdown-induced transcripts and proteomic profiles from cells treated with ICG-001, a β-catenin inhibitor (*104*). This orthogonal validation, spanning both genetic and pharmacological perturbations, and connecting transcriptomic to proteomic responses, suggests that this a *bona fide* biological connection.

This example illustrates how focal graphs can seed systematic pathway mapping efforts. Starting with established pathway components like β-catenin in Wnt signaling, the approach can iteratively expand to explore genetic perturbations, pharmacological modulation, protein-protein interactions, post-translational modifications, genetic dependencies, disease-associated mutations, and cell line profiling that could identify novel targets by which the Wnt pathway might be manipulated for therapeutic benefit.

This approach provides the basis for additional, automated focal graph searches and the foundation for autonomous AI-driven drug discovery, whereby each new connection can serve as a seed for additional focal graph searches. This allows systematic exploration of the pathway’s molecular neighborhood, including cross-validation across multiple experimental approaches, model systems, and data types. Such an iterative strategy could be particularly valuable for identifying new or novel druggable nodes in traditionally challenging pathways in a highly automated, scalable fashion.

## A Framework for Autonomous AI-Driven Drug Discovery

### Wide Applicability of Focal Graphs

As we have shown, focal graphs can be applied to a wide range of drug discovery questions, and can be used to approach individual questions from many perspectives, using multiple data types and model systems. Additional applications in basic research include tool compound identification, model system discovery, metabolomics analysis, natural product target ID, mechanism of resistance analyses, synthetic biology, microbiome analyses, and in studies of rare diseases. They can be applied across multiple stages of the drug discovery pipeline including in the design and characterization of screening libraries, hit to lead optimization, secondary pharmacology (“off-target analyses”), translational medicine, identification of combination therapies, and in the development of companion diagnostics. Further applications include drug repurposing, competitive intelligence, IP diligence & freedom to operate analyses, real world data analysis, veterinary medicine, quality assessment for cell-based therapies, and more. Focal graphs may have applicability across other industries as well, including agriculture, food science, environmental science, cosmetics and personal care, nutraceuticals, forensic science, biodefense and national security, waste management, and bioremediation.

Focal graph approaches have a long list of attributes that make them uniquely powerful and flexible, including transparency, concision, robustness to noise, compatibility with sparse data sets, the capacity to integrate algorithmic approaches including ML-based predictions, and the ability to mine extremely diverse sources of experimental data. These qualities should make them excellent agents for use in an LLM retrieval augmented generation (RAG) application, with focal graph searches being analogous (and complementary) to the internet searches commonly used in RAG-based applications such as Perplexity (*34*).

### Novelty and Bias

The knowledge graphs from which focal graphs are derived have biases that reflect their underlying data sources and the methods of their construction, so it is fair to ask what kinds of bias they may contain. Additionally, much of their information content may have been reported elsewhere, even if only in a database or a supplementary table, so likewise it is fair to ask to what extent they can enable novel discoveries.

Firstly, inclusion of less biased data sources, such as multi-omics data, should lead to less biased results. Secondly, it is important to note that when large scale data sources are reported in the scientific literature, only a small fraction of the data they generate is discussed explicitly in the text. The excess data points - which typically outnumber the text-mentioned data points by many orders of magnitude - are relegated to supplementary files or databases where their retrieval, analysis, and integration face the many hurdles we’ve already discussed. The result is that much of the data - that in theory has been reported - is in practice entirely unknown to the vast majority of researchers.

Finally, connecting dots across many disparate sources is a non-trivial but valuable exercise, which can lead to novel, potentially meaningful connections, as we have seen in many of the drug discovery examples presented here. It’s impractical to ask a human to identify the hypothesis best supported by the data stored in the estimated ∼6,000 existing biological databases,3 or some appreciable fraction thereof, but this is precisely the kind of task that, in principle, can be easily accomplished by a focal graph analysis. In fact, judging by the focal graph results presented so far, such a query is likely to yield an abundance of interesting and novel results.

Furthermore, it would be incorrect to assume that a focal graph analysis is unlikely to bear fruit when applied to obscure or understudied areas, such as rare diseases or novel chemistry. Much is conserved in evolution, and so a gene signature from a rare disease may closely resemble a well-studied biological process, or the knockdown of a well-understood gene in a large-scale omics experiment, yielding clues to its mechanism and suggesting possible therapeutic strategies. Novel chemistry can be mapped to the nearest informative chemical space. Even if the chemical similarity of its neighbors is only modest, integration with other data types may reveal coherent biology that is consistent with the known biological effects of the investigational compound. Alternatively, compound-derived biological profiles, such as multi-omics or cell line proliferation panels, can be used to glean insights into the compound’s target or mechanism independent of its chemical structure.

In summary, although there is certainly bias in knowledge graphs, and by extension, focal graphs, they can nevertheless provide many opportunities for novel discovery; revealing what our “data knows” that human researchers may not.

### Validation

Focal graphs are used here as *information retrieval systems*, much like internet search engines, that can surface information present in large knowledge graphs, and rank what they find by a measure of relevance. Formally speaking, they are not predictive models, although they can be informally used as such, and they could potentially be used to build true predictive models that could then be validated in the usual ways. Because the supporting data is conveyed in a transparent manner, their results can be evaluated in tremendous detail, and the confidence of their conclusions can be determined by domain experts.

Ultimately, validation of hypotheses generated by focal graphs comes down to expert evaluation of the data returned - its reproducibility, its quality, and its quantity - as well as the details of the sources and the methods used to generate it. Science is complex, and the details matter: Specific data points may support one conclusion but not another, and a conclusion that depends on a weak link may not be worth pursuing. Focal graph systems can mine vast amounts of data, surfacing what is most worth considering, generating rich data packages along the way, but in the end it is up to domain experts to judge the level of validation of any given hypothesis.

### Implementation of an Autonomous Target Discovery System

We have built a focal graph-based RAG (FG-RAG) system and have conducted a preliminary analysis of its capabilities. The following prompt was used as a test case: “*Please plan and execute a research program to identify a novel oncology target in the Wnt pathway.*” An excerpted transcript of the response can be found in Supplementary Fig. S2. The FG-RAG proceeds to lay out a plan similar to the focal graph approach described previously, choosing to start with a β-catenin perturb seq profile as a first search, after which it nominates the EIF2 complex as an initial Wnt pathway target candidate, noting, “… *Several components of the eIF2 (eukaryotic initiation factor 2) complex appear in the top Perturb Seq results: [EIF2S1], [EIF2B2], [EIF2B3], [EIF2B4], and [EIF2B5]. This suggests a potential link between protein synthesis regulation and Wnt signaling.* …” The FG-RAG continues its investigation, following up with 3 additional focal graph searches before summarizing its findings and proposing 3 potentially novel Wnt pathway targets: the eIF2 complex, OXA1L (a mitochondrial inner membrane protein), and SSB (Sjogren Syndrome Antigen B).

After this short initial investigation, which was run as a single process in less than 10 minutes, the FG-RAG has the capability to “decide” (or be manually instructed) to continue its perturb-seq strategy with other Wnt pathway members, and/or move on to other focal graph search approaches using a variety of modalities, model systems, data types, etc. Thus the degree of autonomy can be adjusted by the user.

In principle, systems like this can autonomously design and execute research strategies of arbitrary complexity and continue their investigation indefinitely, amassing more and more validation as they go, and increasing the quality, not just the quantity, of the targets ultimately proposed. The extremely modest computational footprint means that truly massive runs of such a system are inherently feasible, and could result in the discovery of highly novel targets with vast amounts of experimental data to support them.

However, in practice, there are notable limitations. For example, the system must be provided with a kernel of potentially successful strategies from which it can generalize, and the results must be evaluated by human experts to ensure the methods and data support the conclusions, an exercise which is made possible by the system’s inherent transparency. While the system can be allowed to run autonomously at a potentially massive scale, the degree of *productive* autonomy will depend on the aforementioned factors, as well as others, constituting a rich area for future studies.

### Towards More Powerful Autonomous Systems

Experimentally supported targets uncovered by focal graphs can be fed into other computational (*105*) or experimental (*106*) approaches, in a unidirectional or iterative way, to design novel therapeutics. These could be small molecules or other modalities, as determined by the scientific, health, and commercial considerations of the project.

The ability of FG-RAG systems to plan, execute, and interpret searches allows the creation of highly elaborate workflows, potentially involving iteration, multiple agent types, hierarchies, heuristics, and other complex decision making structures (*107*, *108*). While individual components of drug discovery have been automated previously (*24*, *109*, *110*), the combination of focal graphs and LLMs allows the autonomous execution of extremely wide ranging research programs, while maintaining full transparency of their scientific rationale, research methods, and the data that supports their findings.

The potential scope of such a global, autonomous research effort is illustrated in Figure 6. The vast space of large-scale biomedical data sets can be explored systematically search by search and study by study, with a tremendous amount and diversity of data being brought to bear on any given research question. While a compelling answer cannot be guaranteed, the system can surface the answer best supported by available data, with subject area experts being the final arbiters of the assembled evidence and conclusions. The system can be further scaled to consider many research questions in parallel, highlighting cases where the data provides a compelling answer that appears to represent a significant scientific or medical breakthrough.

**Figure 6.**
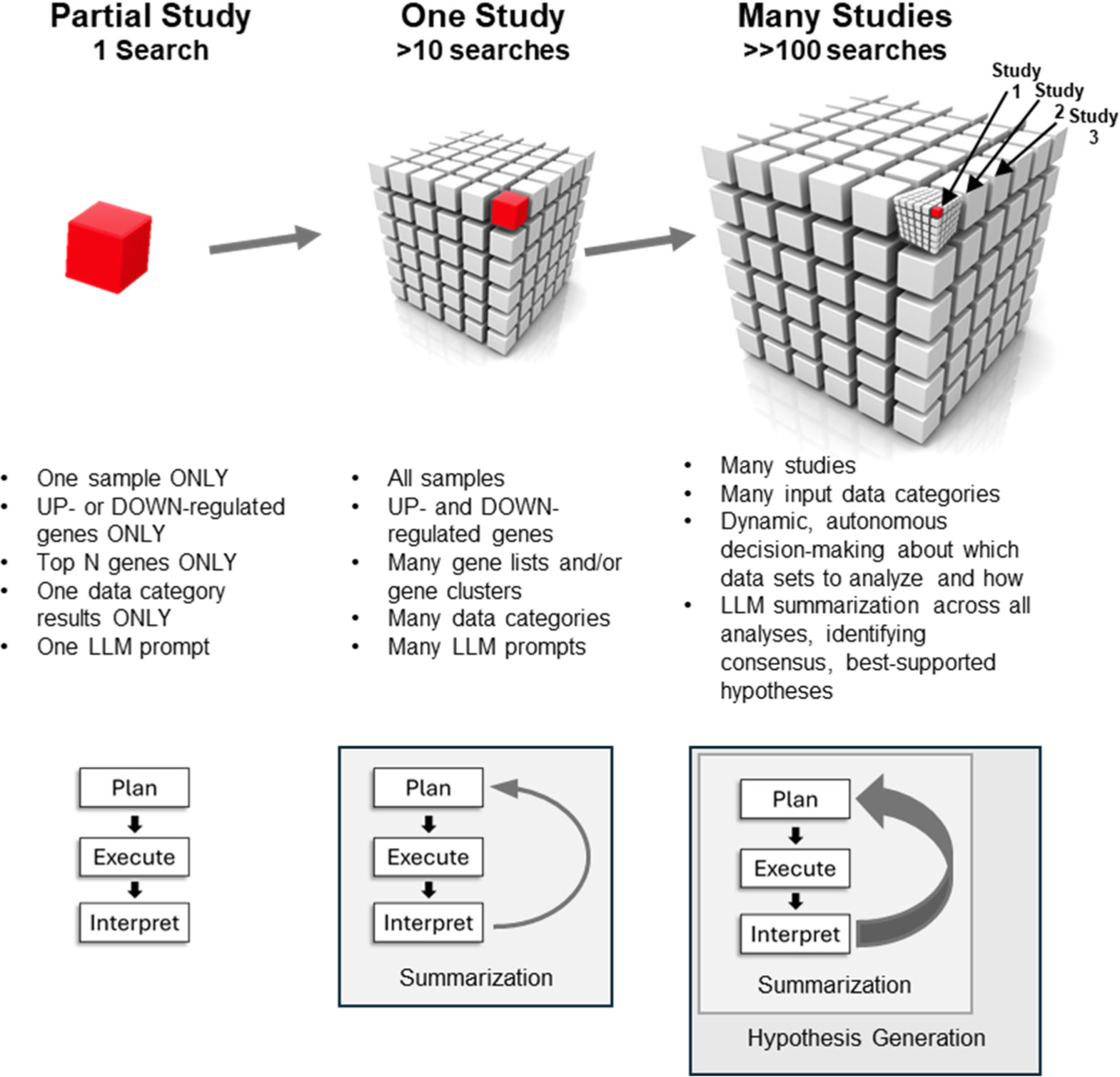
Autonomous planning and execution of research programs at massive scales. The scale of an autonomous RAG-LLM research campaign could be vast, including systematic focal graph searches across large numbers of studies containing many modalities, model systems, data types, and approaches.

A gap analysis could lead to the identification of data sets whose incorporation would maximally increase the information content or novelty of future focal graph search results. Where existing data is insufficient, large-scale data generation campaigns could be launched. Combining an autonomous focal graph approach with a sufficiently powerful robotics-driven system could further increase its power, allowing the automated, targeted generation of key data points that would enable further progress when a purely computational approach would otherwise reach its limit.

## Future Perspective

We have shown here that focal graph approaches are capable of generating novel, data-driven hypotheses from large scale data sets. Although the examples presented are limited, and would admittedly only represent the first steps in a real-world drug discovery program, they nevertheless demonstrate discrete, verifiable progress - achieved by a highly scalable, transparent method that is capable of functioning with a high level of autonomy.

Furthermore, despite the cursory nature of these case studies, we were nevertheless able to uncover a number of interesting and plausible hypotheses backed by specific experimental data. There are an exceptionally wide range of potential focal graph applications, many of which will be the subject of future research reports. These projects can potentially be executed in tremendous depth, including across a breathtaking number of data types, modalities, and model systems. More research is needed to understand the best ways to create and utilize focal graphs, including how to optimally employ them as agents in LLM and RAG frameworks, potentially working side-by-side with other kinds of computational tools and even robotic systems.

It is likely that the answers to many questions of high scientific and medical importance remain hidden in the vast amounts of biomedical data that are largely inaccessible by current methods. This is due to a long list of reasons, including their dispersion across data repositories, their diversity, complexity, variable signal-to-noise, sparseness, scale, and inconsistent annotation, among others. We believe that the focal graph-based approaches described here can make a significant contribution. When combined with LLMs and other tools, focal graphs could transform our rich but underutilized data inheritance into a deeper understanding of the mechanisms underlying disease and into breakthrough therapies that could improve the human condition.

For too long, drug discovery efforts have been ineffective at using the vast amount of data produced by the global biomedical community. Each project generates and regenerates data anew, costing an unconscionable amount of time, resources, and ultimately, lives. It is past time that this effort became cumulative, with each new project truly standing on the shoulders of the giants that came before it. With the powerful tools we now have in hand - and with more appearing on the horizon - we may finally be on the verge of breaking through this tragic impasse. By learning from our historical data, rather than being doomed to forever repeat it, we may be able to make huge strides in our understanding of disease and in our relentless quest for cures.

## Supporting information

Selinger et al Supplement

## Acknowledgments

We extend our sincere gratitude to Dr. George Church and Andy Palmer for their insightful feedback and unwavering support for this research.

This work was supported by the Zoll Foundation award (to O.L) and by the NIAID (NIH) under Award Number R21AI196527 (to O.L). The content is solely the responsibility of the authors and does not necessarily represent the official views of the NIH.

## Author Contributions

D.W.S.: Conceptualization, Methodology, Data Analysis, Formal Analysis, Project Administration, Supervision, Investigation, Funding Acquisition, Visualization, Writing - Review & Editing, Writing - Original Draft

T.R.W.: Software, Methodology, Formal Analysis, Visualization, Writing - Review & Editing

E.S.: Conceptualization, Data Analysis, Project administration, Visualization, Writing - Review & Editing

E.M.K.: Conceptualization, Data Analysis, Visualization, Writing - Review & Editing

J.G.: Conceptualization, Data Analysis, Visualization, Writing - Review & Editing

O.L.: Conceptualization, Data Analysis, Project Administration, Visualization, Writing - Review & Editing

## Declaration of Interests

The authors declare the following interests in Plex Research, Inc.: D.W.S. is a founder, employee, shareholder, and member of the board of directors; T.R.W. and J.G. are employees and option holders; E.S. and E.M.K. are consultants; O.L. is a consultant and investor.

## Declaration of Generative AI and AI-assisted Technologies in the Writing Process

During the preparation of this work, the authors used Claude 3.5 Sonnet (https://claude.ai) and Perplexity (https://www.perplexity.ai) in order to suggest phrasing and composition and to extract text from images. After using these tools/services, the authors reviewed and edited the content as needed and take full responsibility for the content of the publication.

## Notes

### Competing Interest Statement

All authors are employees or consultants of Plex Research, Inc. and may have real or optional ownership therein.

### Summary of Updates

- A new quantitative analysis of compound target prediction accuracy of an LLM both with and without a focal graph search - Additional details on the methodology used to generate focal graphs - An extended discussion of the nature and extent of the system's autonomy - References to newly published and submitted studies that utilized and/or validated focal graph results

