## Supplementary material for "A Framework for Autonomous AI-Driven Drug Discovery": Selinger et al Supplement

**This PDF file includes:**

Supplementary Methods
Supplementary Figs. S1 and S2
Supplementary References

**Other Supplementary Materials for this manuscript include the following:**

Supplementary File: Selinger_etal_Autonomous_AI_Tables S1-S6.xlsx
Including:
Supplementary Table S1: S1_Compound input table
Supplementary Table S2: S2_Output_LLM-only
Supplementary Table S3: S3_Output_Focal Graph-LLM_ecfp
Supplementary Table S4: S4_Evaluation_detailed_results
Supplementary Table S4: S5_Output_Focal Graph-LLM_sim
Supplementary Table S6: S6_Secondary analysis

**Supplementary Methods**

#### Focal graph construction

The focal graph construction uses degree-based centrality with category-specific edge weights (edge_score = 1 + boost), accumulated bidirectionally during compilation. Evidence sources are aggregated by summing edge weights across all connections between node pairs, with category-specific upper range compression and hub filtering applied to prevent ubiquitous entities from dominating. Supporting evidence strength is incorporated into the LLM interpretation through the accumulated edge weight passed to the AI agent, which receives neighbor lists sorted by aggregated weight and uses this quantitative signal to assess relationship importance.

Graph search expansion retrieves neighbors for a given starting node in category-aware hops: starting from query nodes (e.g., compounds from similarity search), the system fetches weighted neighbors by category (targets, diseases, pathways), applies hub filtering via weight cutoffs, aggregates edge weights across all source nodes, and returns top-N neighbors sorted by accumulated weight. Multi-hop expansion repeats this process using the previous hop's results as new source nodes, building outward from the query while maintaining category boundaries - each hop's neighbors are pooled by category, filtered against degree centrality thresholds, and ranked by the sum of their edge weights (where edge weight = 1 + boost per connection). The expansion continues for the specified number of hops (typically 1-3), with each successive hop potentially introducing thousands of new nodes, though neighbor limits and weight cutoffs prevent exponential explosion by prioritizing high-centrality connections and recent relationships.

#### FG-LLM Quantitative Benchmarking

##### 1. Overview and Rationale

We designed a quantitative benchmark for chemical structure-based target identification to compare the accuracy of a focal graph-enabled LLM (FG-LLM) to the same LLM lacking focal graph search. To increase the benchmark's relevance for real-world applications where the targets of compounds are unknown, we created a modified reference set, based on the MoA Box , where each compound was modified by the addition of a single fluorine atom. This change is small enough that it is unlikely to change the compounds' targets but it increases the system's reliance on chemical structure-based inferences rather than on trivial target lookups, which would not be possible in any case when analyzing novel compounds.

##### 2. Source Dataset: The MoA Box Collection

The parent compound library was drawn from the MoA Box, a curated collection of 4,185 small molecules with expert-assigned target annotations and mechanism-of-action classifications (1). The  Box was selected because its annotations represent high-confidence, manually curated assignments across a broad chemical and biological space, providing a reliable gold standard against which FG-LLM predictions could be assessed. Each compound in the collection is associated with one or more primary target gene symbols and mechanism-of-action descriptions, enabling automated, quantitative matching to gene symbols, as well as more thorough, manual evaluation based on full MoA annotations.

#### 3. *In Silico* Generation of Monofluorinated Derivatives

For each compound in the MoA Box, a set of candidate monofluorinated derivatives was generated computationally by systematic enumeration of viable C-H to C-F substitutions using the RDKit extension in KNIME software (2, 3). Eligible substitution sites were defined using SMARTS-based substructure queries and restricted to aromatic carbons and non-carbonyl aliphatic carbons, consistent with positions commonly targeted in medicinal chemistry fluorination strategies and less likely to substantially disrupt compound stability or binding geometry (4–6). Carbonyl-adjacent positions were excluded to avoid generating acyl fluorides or other reactive or metabolically labile species.

From the enumerated candidate structures for each parent compound, the single derivative exhibiting the highest structural similarity to its parent was selected. Molecular similarity was calculated using the Tanimoto coefficient applied to binary extended connectivity fingerprints with diameter 2 (ECFP2) (7) (see Supplementary Table S1)

Following derivative selection, each candidate structure was screened against the knowledge graph to confirm the absence of any existing annotations, bioactivity data, or structural records within the FG-LLM system being tested. Compounds with any pre-existing representation in the graph were excluded from the test set. This filtering step ensures that any FG-LLM predictions for these compounds arose from inference based on structural similarity and focal graph analysis, rather than direct lookup of the compounds themselves.

#### 4. FG-LLM Workflow Applied to the Test Set

A random subset of 500 compounds was drawn from the filtered set of fluorinated derivatives for benchmarking. Randomization was performed without stratification to ensure an unbiased sample of chemical and target diversity.

For each of the 500 compounds, the FG-LLM workflow was executed using KNIME software as follows. First, a focal graph (FG) search was executed for each compound using consistent, pre-specified parameters across all compounds. Specifically, molecular similarity expansion for the knowledge graph query used the Tanimoto coefficient applied to binary extended connectivity fingerprints with diameter 4 (ECFP4) (7). The FG step retrieves structurally similar compounds and their associated bioactivity annotations from the knowledge graph, returning a ranked set of neighbors with associated target and MoA information. Second, these FG results were provided as context to the large language model (Anthropic Claude Sonnet 4.5), along with a standardized prompt instructing the model to use the focal graph query results to infer the most likely mechanisms of action, target gene symbols, and other relevant properties of the query molecule. The prompt was held constant across all compounds to eliminate prompt variability as a source of performance differences.

In parallel, the same LLM was prompted to perform identical predictions for each of the 500 compounds without access to focal graph results. The prompt used for this LLM-only condition was generated from the same master template, differing from the FG-LLM prompt only in the substitution of template variables for compound identification, referencing the focal graph results, and fingerprint-specific similarity values. Specifically, the LLM-only prompt instructed the model to draw on all information available to it about the compound indicated by its SMILES, rather than directing it to prioritize focal graph search results. All other instructions, output schema, controlled vocabularies, and evidence hierarchy guidance were identical between conditions. Refer to the Prompt Template section of this document for more information.

#### 6. Performance Evaluation Against MoA Box Annotations

Predicted targets were evaluated against MoA Box annotations for the pre-fluorinated parent compound of each test molecule (see Supplementary Tables S2-S4). The primary evaluation metric was exact gene symbol match: a prediction was counted as a hit if any MoA Box-annotated target gene symbol for the parent compound appeared within the predicted top 1 or top 10 targets. Hit rates were calculated for both top 1 and top 10 thresholds.

#### 7. Secondary Analysis of Non-Matching Compounds

The 36 compounds for which FG-LLM failed to predict any MoA Box-annotated target within the top-10 results were subjected to a secondary analysis. These compounds were reprocessed using an alternative similarity algorithm, namely the Indigo Toolkit sim fingerprint, providing a different retrieval strategy. This second-pass FG-LLM run recovered a matching target for 8 of the 36 compounds (22.2%), reducing the unresolved set to 28 compounds (see Supplementary Tables S5).

The remaining 28 compounds were subjected to manual review to characterize the nature of the apparent discrepancies. We assessed whether FG-LLM predictions reflected biologically meaningful relationships to MoA Box annotations that were not captured by exact gene symbol matching (see Supplementary Tables S6). The following categories of partial or indirect concordance were identified:

- Five compounds: FG-LLM-inferred targets matched the descriptive mechanism of action recorded in MoA Box-annotated "moa" field, but failed exact gene symbol matching due to incomplete or non-standard MoA Box annotations (e.g., MoA Box annotating targets or sets of targets in the "moa" description without full annotation of relevant target gene products in "gene_symbols" fields).
- Three compounds: FG-LLM predicted a different member of the same protein family as the annotated target, consistent with cross-selectivity patterns commonly observed for compounds targeting multi-member protein families.
- Five compounds: FG-LLM predicted targets belonging to the same target class as the annotated target, or targets sharing identical endogenous ligands (e.g. glutamate receptors), indicating likely pharmacological relevance despite inexact gene-level correspondence.
- Six compounds: FG-LLM predicted targets within the same biological pathway as the annotated target, which may reflect a limitation in precision.
- One compound, a phorbol ester (MOABOX:5922), was annotated by our FG-LLMLLM as a putative protein kinase C activator, while the parent compound is annotated with the target TRPV4 in the MOA Box. Given that phorbol esters are well-known, canonical PKC activators, there is high likelihood the FG-LLM annotation is correct (8).
- Eight compounds: No discernible concordance between FG-LLM predictions and MOA Box annotations under any of the above criteria.

### Prompt Template

#### Overview

Prompts for the quantitative benchmarking analysis were derived from a master template (below). The template was designed to create essentially equivalent prompts while still allowing the provision of focal graph-specific parameters to the FG-LLM. The values of the parameters included as variables (red) can be found in the table following the template.

#### Master Prompt Template

You are a scientific model trained to infer compound mechanisms of action (MOA) and molecular targets using the ChEMBL schema and vocabulary.
Carefully review **{{DATA_SOURCE}} {{COMPOUND_REFERENCE}}**, prioritizing **{{PRIORITY_CATEGORIES}}**.

Also, prioritize evidence quality over quantity.

- Start with direct compound-target relationships and their evidence quality
- Use other types of information (publications, patents, diseases, protein families) as supporting context
- Do not let high counts in indirect categories (publications, diseases) override direct mechanistic data
- If similar compounds have curated MOA annotations, these should be primary hypotheses

**Evidence Quality Hierarchy (highest to lowest priority):**

- Direct experimental binding/activity data with quantitative measurements
- Curated mechanism of action annotations from specialized databases (ChEMBL MOA, DrugBank, etc.). These represent expert curation and should be prioritized
- Direct compound-target relationships from binding databases (BindingDB, ChEMBL) with activity data
- Indirect relationships through highly similar compounds**{{SIMILARITY_THRESHOLD_EVIDENCE}}** with direct data
- Literature co-mentions and patent associations
- Protein family/complex associations without direct binding data

Based on your analysis, infer the most likely:

- Mechanism of Action
- Action Type
- Target(s) Name(s)
- Target(s) Symbol(s)
- Target Type
- Confidence Level

**Field definitions:**
**Target Name:** Official name of target(s) or target(s) family (e.g., "Dual specificity mitogen-activated protein kinase kinase; MEK1/2").
**Target Symbol:** Official gene symbol(s) for the primary target(s) (HGNC symbol for human by default; for non-human targets, use the appropriate official species symbol or standard gene symbol).

- If multiple targets are involved, list them in matching order separated by semicolons.
- If the symbol cannot be determined, use "UNKNOWN".
- Do NOT use slashes ("/"), or any other characters to separate target symbols. If multiple family members are targets, list them separately, separated by semicolons.

Many bioactive compounds have multiple targets, or even many targets. When similar compounds show evidence for multiple targets, report ALL targets with substantial evidence rather than selecting a single "primary" target. Use the "Target Name" and "Target Symbol" fields to list multiple targets separated by semicolons, ordered by evidence strength.

Assume human targets by default unless there is a clear indication of a different species.

**Controlled vocabularies:**
List of acceptable Action Type terms (ChEMBL):
ACTIVATOR, AGONIST, ALLOSTERIC ANTAGONIST, ANTAGONIST, ANTISENSE INHIBITOR, BINDING AGENT, BLOCKER, CHELATING AGENT, CROSS-LINKING AGENT, DEGRADER, DISRUPTING AGENT, EXOGENOUS GENE, EXOGENOUS PROTEIN, GENE EDITING NEGATIVE MODULATOR, HYDROLYTIC ENZYME, INHIBITOR, INVERSE AGONIST, METHYLATING AGENT, MODULATOR, NEGATIVE ALLOSTERIC MODULATOR, NEGATIVE MODULATOR, OPENER, OTHER, OXIDATIVE ENZYME, PARTIAL AGONIST, POSITIVE ALLOSTERIC MODULATOR, POSITIVE MODULATOR, PROTEOLYTIC ENZYME, REDUCING AGENT, RELEASING AGENT, RNAI INHIBITOR, SEQUESTERING AGENT, STABILISER, SUBSTRATE, VACCINE ANTIGEN

List of acceptable Target Type terms (ChEMBL):
ADMET, CELL-LINE, CHIMERIC PROTEIN, MACROMOLECULE, NO TARGET, NON-MOLECULAR, NUCLEIC-ACID, ORGANISM, PROTEIN COMPLEX, PROTEIN COMPLEX GROUP, PROTEIN FAMILY, PROTEIN-PROTEIN INTERACTION, SELECTIVITY GROUP, SINGLE PROTEIN, SMALL MOLECULE, SUBCELLULAR, TISSUE, UNCHECKED, UNKNOWN

**Structural Similarity Inference Rules:**
When inferring mechanism of action and targets**{{CONTEXT_PHRASE}}**, evaluate structural similarity quantitatively rather than qualitatively - compounds with **{{SIMILARITY_THRESHOLD_HIGH}}** should be considered structurally equivalent for biological activity predictions. Geometric isomers, stereoisomers, or compounds differing only in minor structural features (salts, hydrates, tautomers) with high similarity**{{SIMILARITY_SCORES_TEXT}}** warrant high confidence inferences when experimental data exists for the similar compound. Only reduce confidence for "similar compound" data when similarity**{{SIMILARITY_SCORES_TEXT}}** is **{{SIMILARITY_THRESHOLD_MODERATE}}** or when significant structural differences exist that could impact the proposed mechanism. The phrase "data derived from a similar compound" should not automatically lower confidence - instead, assess whether the structural differences are mechanistically relevant to the target interaction.

Format the output as a JSON.
Output JSON:
```
{
  "Mechanism of Action": "",
  "Action Type": "",
  "Target Name": "",
  "Target Symbol": "",
  "Target Type": "",
  "Confidence Level": ""
}
```

#### Template Variables

LLM-only Configuration

| Variable | Value |
|---|---|
| {{DATA_SOURCE}} | all information available to you |
| {{COMPOUND_REFERENCE}} | about the compound indicated by the SMILES `[SMILES]` |
| {{PRIORITY_CATEGORIES}} | information about Targets, Patents, and Publications |
| {{CONTEXT_PHRASE}} | (omit - empty string) |
| {{SIMILARITY_THRESHOLD_EVIDENCE}} | (omit - empty string) |
| {{SIMILARITY_THRESHOLD_HIGH}} | high similarity |
| {{SIMILARITY_SCORES_TEXT}} | (omit - empty string) |
| {{SIMILARITY_THRESHOLD_MODERATE}} | moderate |

Focal graph+LLM Configurations

| Variable | Value (when using ECFP4 fingerprint) (default) | Value (when using Indigo Toolkit sim fingerprint) |
|---|---|---|
| {{DATA_SOURCE}} | these focal graph results | these focal graph results |
| {{COMPOUND_REFERENCE}} | (omit - empty string) | (omit - empty string) |
| {{PRIORITY_CATEGORIES}} | the Targets, Patents, and Publications categories | the Targets, Patents, and Publications categories |
| {{CONTEXT_PHRASE}} | from focal graph search results | from focal graph search results |
| {{SIMILARITY_THRESHOLD_EVIDENCE}} | (similarity ≥ 0.5) | (similarity ≥ 0.8) |
| {{SIMILARITY_THRESHOLD_HIGH}} | similarity scores ≥0.55 or perfect 1.000 scores | similarity scores ≥0.9 or perfect 1.000 scores |
| {{SIMILARITY_SCORES_TEXT}} | scores | scores |
| {{SIMILARITY_THRESHOLD_MODERATE}} | moderate (0.3-0.5) | moderate (0.7-0.89) |

#### Fig. S1. LLM Comparison

**Query Items:** Apoe, Lyz2, Spp1, Lgals3, Postn, H19, Col3a1, Fn1, Bgn, Ctsb, Ctss, Vim, Tmsb10, Mt1, Ftl1, Tyrobp, Col1a1, Ctsl, S100a4, Sln, Myl4, H2-D1, Cyba, Cd74, Mgp, Cfd, Eef1a1, Grn, Col1a2, S100a6, C1qa, Fbxl22, Tuba1b, Ctsz, C1qc, Ckb, Mustn1, H2-Aa, Lgmn, Gpx1, C1qb, Mt2, Ctsk, H2-Ab1, Rpl3, Crip1, Fcer1g, Selenop, Fabp5, Anxa2

**Prompt:** This search was run with a set of genes that are up-regulated in a disease. Based on the profiling results, what disease do you think it might be? Please provide any evidence you found to support your hypothesis.

**Focal Graph - LLM**

To answer this question, I'll examine the profiling results, particularly focusing on the RNA Profile, Proteomic Profile, and Biomarker categories, as these are most likely to provide insights into disease-related gene expression patterns:

"gxa:E-GEOD-1025 Hindlimb muscle, comparison of wild type and mdx mice, 7 to 112 Day (Porter lab)" "geo:GDS4200 Cmah knockout effect on mdx model of Duchenne Muscular Dystrophy (DMD): skeletal muscle" "geo:GDS703 Dystrophin-deficient mdx extraocular and leg muscle" "geo:GDS3398 Utrophin overexpression effect on dystrophin-deficient mdx skeletal muscles" "gxa:E-GEOD-466 mRNA expression in regenerated mdx mouse skeletal muscle" "geo:GDS236 Dystrophin-deficient mdx muscle regeneration"

Based on these search results, **the disease that this gene set is most likely associated with is Duchenne Muscular Dystrophy (DMD).** Here's the evidence to support this hypothesis:

***1. Multiple RNA profiles in the search results specifically mention "mdx" mice, which are a well-established animal model for Duchenne Muscular Dystrophy.*** For example: Hindlimb muscle comparison of wild type and mdx mice, Dystrophin-deficient mdx extraocular and leg muscle, mRNA expression in regenerated mdx mouse skeletal muscle

***2. One of the RNA profiles explicitly mentions DMD:*** Cmah knockout effect on mdx model of Duchenne Muscular Dystrophy (DMD): skeletal muscle

***3. The search results include several references to muscle-related studies, which is consistent with DMD being a muscular disorder:*** Utrophin overexpression effect on dystrophin-deficient mdx skeletal muscles, Dystrophin-deficient mdx muscle regeneration

***4. The presence of studies involving muscle regeneration and comparisons between wild-type and mdx mice suggest that the gene set is related to a progressive muscle disorder, which is characteristic of DMD.***

While this **evidence strongly points to Duchenne Muscular Dystrophy,** it's important to note that this is a hypothesis based on the available search results. Other muscular dystrophies or related disorders could potentially show similar gene expression patterns. To confirm this hypothesis, further analysis of the specific genes in the query set and their known associations with DMD would be beneficial.

**Claude 3.5 Sonnet (Anthropic)**

*Based on the list of up-regulated genes provided, I believe **this gene expression profile might be associated with cardiac fibrosis or heart failure.** Here's the evidence to support this hypothesis:*

*1. Extracellular matrix (ECM) remodeling genes: Several genes in the list are associated with ECM production and remodeling, which is a hallmark of cardiac fibrosis: Col1a1, Col1a2, Col3a1 (collagen genes), Fn1 (fibronectin), Postn (periostin), Bgn (biglycan), Mgp (matrix Gla protein)*

*2. Fibroblast activation markers: S100a4 (also known as FSP1, fibroblast-specific protein 1), Vim (vimentin)*

*3. Inflammatory and immune response genes: Chronic inflammation is often associated with heart failure and fibrosis:*

*Cd74, H2-Aa, H2-Ab1, H2-D1 (MHC class II-related genes); C1qa, C1qb, C1qc (complement system components), Tyrobp, Fcer1g (immune cell signaling)*

*4. Cardiac stress markers: Apoe (apolipoprotein E, associated with cardiovascular disease), Spp1 (osteopontin, involved in cardiac remodeling), Myl4 (atrial light chain-1, associated with atrial fibrillation)*

*5. Oxidative stress and cellular stress response: Mt1, Mt2 (metallothioneins, involved in oxidative stress response), Gpx1 (glutathione peroxidase 1, antioxidant enzyme)*

*6. Proteases and cathepsins: These are involved in ECM degradation and remodeling: Ctsb, Ctsk, Ctsl, Ctsz (cathepsins)*

*7. Other relevant genes: Lgals3 (galectin-3, a biomarker for heart failure), Sln (sarcolipin, regulates calcium cycling in cardiomyocytes)*

*The combination of ECM remodeling genes, inflammatory markers, and cardiac stress-related genes strongly suggests a fibrotic cardiac condition. The presence of genes like Apoe, Lgals3, and Spp1 further supports the hypothesis of heart failure or cardiac fibrosis.*

*It's important to note that while **this gene expression profile is highly suggestive of cardiac fibrosis or heart failure**, additional clinical and experimental data would be needed to confirm the diagnosis. The profile could also be consistent with fibrosis in other organs, so cardiac-specific markers and clinical context would be crucial for a definitive diagnosis.*

**ChatGPT4o (OpenAI)**

*The genes listed are involved in various biological processes, particularly in inflammation, fibrosis, immune response, and extracellular matrix remodeling. The upregulation of these genes is characteristic of several pathological conditions, often related to chronic inflammation, fibrosis, and immune system activation.*

*Key Genes and Their Roles*

*1. Apoe (Apolipoprotein E): Associated with Alzheimer's disease, cardiovascular disease, and inflammation. Spp1 (Osteopontin): Involved in inflammation and fibrosis, commonly elevated in chronic inflammatory diseases, cancer, and autoimmune diseases. Lgals3 (Galectin-3): Involved in fibrosis, inflammation, and cancer. Col1a1, Col3a1 (Collagens): Major components of the extracellular matrix, typically upregulated in fibrotic conditions (e.g., liver fibrosis, pulmonary fibrosis). Fn1 (Fibronectin): Another extracellular matrix protein upregulated in tissue injury and fibrosis. Myl4 (Myosin light chain 4): Typically involved in muscle function, but its upregulation is seen in cardiac diseases, especially those related to heart failure or fibrosis. Cd74, H2-D1, H2-Ab1: MHC class II genes, indicating immune system activation, potentially in autoimmune diseases or chronic inflammatory responses. C1qa, C1qb, C1qc: Components of the complement system, often associated with autoimmune diseases like systemic lupus erythematosus (SLE). S100a4, S100a6: S100 proteins are involved in inflammation, cancer, and fibrotic diseases. Ftl1 (Ferritin light chain): May indicate an oxidative stress response, commonly associated with neurodegenerative diseases like Alzheimer's. Crip1 (Cysteine-rich protein 1): Often involved in immune and stress response.*

***Potential Disease Hypothesis***

***The gene signature suggests involvement in fibrosis-related diseases, chronic inflammatory conditions, and potentially autoimmune diseases.*** *The upregulation of extracellular matrix genes (Collagens, Fibronectin), fibrosis markers (Spp1, Lgals3), immune system activation (Cd74, MHC class II genes), and complement activation (C1q genes) points towards diseases like:- Idiopathic Pulmonary Fibrosis (IPF): Given the strong extracellular matrix and immune involvement.- Systemic Lupus Erythematosus (SLE): Due to immune activation and complement involvement.- Alzheimer's Disease or Neurodegenerative Diseases: Apoe, S100 proteins, and ferritin light chain are implicated in neurodegenerative diseases.- Cardiac Fibrosis or Heart Failure: Myl4 and other fibrosis-related genes are upregulated in heart conditions.*

*Supporting Evidence:*

*-Fibrosis-related genes(Collagens, Fn1, Lgals3) suggest diseases with significant tissue remodeling and scarring.*

*- Immune activation(Cd74, C1qa, H2-D1) supports the involvement of immune-mediated or autoimmune diseases.*

*- Apoe and S100 proteins link to neurodegeneration and chronic inflammation.*

**Gemini 1.5 Pro (Google)**

*The provided gene list* ***strongly suggests fibrosis as the disease.*** *Here's why:*

*Evidence:*

*- Extracellular Matrix (ECM) Remodeling: A significant number of the genes are directly involved in the production and remodeling of the ECM, a hallmark of fibrosis. These include: Col1a1, Col1a2, Col3a1: Collagen genes, the main structural proteins of the ECM. Upregulation points to excessive collagen deposition. Fn1: Fibronectin, another crucial ECM glycoprotein involved in cell adhesion and migration. Postn: Periostin, an ECM protein that interacts with collagen and promotes fibrosis. Spp1: Osteopontin, a matricellular protein involved in inflammation and ECM remodeling. Timp1: Tissue inhibitor of metalloproteinase 1, regulates ECM degradation and its upregulation indicates a shift towards ECM deposition.*

*Mmp2, Mmp9: Matrix metalloproteinases, involved in ECM breakdown. While typically associated with ECM degradation, their upregulation can also be found in certain fibrotic conditions, suggesting a dysregulated remodeling process.*

*Inflammation and Immune Response: Fibrosis is often preceded and perpetuated by inflammation. The presence of these genes supports this link:*

*- Lgals3 (Galectin-3): A pro-inflammatory and pro-fibrotic lectin.*

*- Ctsb, Ctsl, Ctsk: Cathepsins, lysosomal proteases involved in inflammation and ECM degradation.*

*- S100a4: A calcium-binding protein implicated in inflammation and fibrosis.*

*- H2-Aa, H2-Ab1, H2-D1: Major histocompatibility complex (MHC) class II genes, indicating immune system activation.*

*- Cd74: MHC class II-associated invariant chain involved in antigen presentation and inflammatory responses.*

*Other Supporting Genes:*

*- Apoe: Apolipoprotein E, linked to lipid metabolism and inflammation. Its upregulation is observed in various fibrotic conditions.*

*- Acta2 (α-SMA): Alpha-smooth muscle actin, a marker of myofibroblast differentiation, the key cell type responsible for fibrosis.*

*- Tgfb1: Transforming growth factor beta 1, a potent pro-fibrotic cytokine. While not in your list directly, its downstream targets are heavily represented.*

*Important Considerations:*

*- Tissue Specificity: Fibrosis can occur in various organs (lung, liver, heart, kidney, etc.). Further investigation is necessary to pinpoint the specific type of fibrosis.*

*- Disease Stage: The gene expression profile can vary depending on the disease's stage (early vs. late).*

*- Confirmatory Testing: This analysis provides a strong hypothesis. Further investigations, including histological examination, clinical evaluation, and potentially more targeted gene expression analysis, are crucial for diagnosis.*

**MedLM-Large (Google)**

*The genes listed are all up-regulated in* ***atherosclerosis.*** *Atherosclerosis is a disease in which plaque builds up inside the arteries. This can lead to a number of serious problems, including heart attacks and strokes. The genes listed are involved in a variety of processes, including inflammation, cell growth, and the formation of new blood vessels.*

**Co-Pilot Pro (Microsoft)**

*Based on the list of up-regulated genes you provided, it appears that these genes are associated with several diseases, but* ***one prominent possibility is Alzheimer's disease.*** *Here are some key points supporting this hypothesis:*

*1. Apoe (Apolipoprotein E): This gene is well-known for its association with Alzheimer's disease. The ApoE4 variant is a major genetic risk factor for late-onset Alzheimer's.*

*2. Spp1 (Secreted Phosphoprotein 1): Also known as osteopontin, this gene has been implicated in neuroinflammation and is up-regulated in Alzheimer's disease.*

*3. Lgals3 (Galectin-3): This gene is involved in inflammation and has been found to be up-regulated in Alzheimer's disease.*

*4. Postn (Periostin): This gene is associated with extracellular matrix remodeling and has been linked to neurodegenerative diseases.*

*5. H19: This long non-coding RNA is involved in various cellular processes and has been found to be dysregulated in Alzheimer's disease.*

*6. Col3a1, Col1a1, Col1a2 (Collagens): These genes are involved in extracellular matrix organization and have been associated with Alzheimer's disease.*

*7. Fn1 (Fibronectin 1): This gene is involved in cell adhesion and migration and has been implicated in Alzheimer's disease.*

*8. Ctsb, Ctsl, Ctsz (Cathepsins): These lysosomal proteases are involved in protein degradation and have been found to be up-regulated in Alzheimer's disease.*

*9. S100a4, S100a6 (S100 Calcium Binding Proteins): These proteins are involved in calcium signaling and have been associated with Alzheimer's disease.*

*10. Tyrobp (TYRO Protein Tyrosine Kinase Binding Protein): This gene is involved in immune response and has been linked to Alzheimer's disease.*

*These genes collectively suggest* ***a strong association with Alzheimer's disease****, particularly due to their roles in neuroinflammation, extracellular matrix remodeling, and immune response.*

**Perplexity Pro (Perplexity)**

*Based on the list of upregulated genes provided,* ***the disease in question is likely Alzheimer's disease (AD)****. Several key pieces of evidence from the search results support this hypothesis:*

*1. Apolipoprotein E (APOE) is prominently featured in the gene list. APOE is one of the strongest genetic risk factors for Alzheimer's disease. The APOE ε4 allele in particular is associated with increased risk and severity of AD.*

*2. Many of the listed genes are associated with microglial activation and neuroinflammation, which are hallmarks of Alzheimer's disease:*

*- Tyrobp, C1qa, C1qb, C1qc, Cd74, and Fcer1g are involved in microglial activation and immune responses in the brain*

*- Lyz2, Spp1, Lgals3, Ctsb, Ctss, and Ctsl are markers of activated or disease-associated microglia (DAM) in AD*

*3. Several genes in the list are part of the "disease-associated microglia" (DAM) signature identified in Alzheimer's disease models:*

*- Apoe, Tyrobp, Ctsd, Cst7, and Spp1 are specifically mentioned as part of the DAM signature in the search results*

*4. The presence of genes related to oxidative stress and redox regulation (e.g., Gpx1, Mt1, Mt2) aligns with the known oxidative stress component of Alzheimer's disease pathology.*

*5. Complement-related genes (C1qa, C1qb, C1qc) are upregulated, which is consistent with the involvement of the complement system in AD pathogenesis.*

*6. Genes involved in lipid metabolism and transport (Apoe, Fabp5) are present, reflecting the altered lipid metabolism observed in AD.*

*7. The upregulation of genes related to phagocytosis and lysosomal function (Ctsb, Ctss, Ctsl, Ctsz, Lgmn) is consistent with the increased phagocytic activity of microglia in AD, particularly around amyloid plaques.*

*While this gene expression profile could potentially be associated with other neurodegenerative or neuroinflammatory conditions, the strong presence of AD-associated genes, particularly APOE and the DAM signature, strongly suggests Alzheimer's disease as the most likely candidate. The combination of microglial activation markers, oxidative stress-related genes, and AD risk factors provides compelling evidence for this conclusion.*

#### Fig. S2. FG-RAG Transcript (Excerpt)

**Human User: Please plan and execute a research program to identify a novel oncology target in the Wnt pathway.**

**AI Guide Agent: Certainly! I'll plan and execute a research program to identify a novel oncology target in the Wnt pathway using focal graphs. We'll follow a step-by-step approach, leveraging the power of focal graph to uncover potential new targets.**

<u>Step 1: Identify known key players in the Wnt pathway</u>
First, let's start with some well-known genes in the Wnt pathway that are associated with cancer. We'll use these as our starting points. Some key players include:
1. CTNNB1 (β-catenin)
2. APC
3. GSK3B
4. AXIN1
5. WNT1

<u>Step 2: Resolve focal graph IDs for Perturb-seq profiles of these genes</u>
Now, let's search for Perturb-seq profiles for each of these genes. We'll do this one at a time to ensure we get the correct profiles.
Resolved IDs: gwps:1937_CTNNB1_P1P2_ENSG00000168036|dn, gwps:1937_CTNNB1_P1P2_ENSG00000168036|up
It seems there are Perturb-seq profiles available for both the up- and down-regulated genes after [CTNNB1] knockdown. We'll analyze both, starting with the up-regulated Perturb-seq profile for [CTNNB1](https://gwps.wi.mit.edu/?chosen_gene=ENSG00000168036 "gwps:1937_CTNNB1_P1P2_ENSG00000168036|up").

<u>Step 3: Perform a focal graph search with the identified [CTNNB1] Perturb-seq profile</u>
Let's run a focal graph search using the up-regulated Perturb-seq profile for [CTNNB1]
**AI Guide Agent: Analyze the results, focusing on the target category. Produce a list of the top 200 targets (or as many as are available if less than 200) from this search.**

**AI Analyst Agent: Certainly! I'll analyze the results of the focal graph search, focusing on the target category, and provide you with a list of the top 200 targets (or as many as are available if less than 200). Based on the results from the focal graph search, I have analyzed the target category and compiled a list of the top 200 targets. Here are the top 200 targets (or as many as were available) from this search, listed by their Entrez Gene IDs:**

1. [IDO1](entrezgene:3417)
2. [MAPK1](entrezgene:5594)
3. [MAPK14](entrezgene:1432)
4. [EPN1](entrezgene:9682)
5. [RAB4A](entrezgene:51135)
6. [THRB](entrezgene:8743)
7. [MTNR1A](entrezgene:4595)
8. [TUG1](entrezgene:7846)
9. [RELA](entrezgene:5970)
10. [CTPS1](entrezgene:1520)
11. [LGALS3](entrezgene:3956)
12. [IKBKB](entrezgene:3551)
13. [FZD5](entrezgene:2534)
14. [TXNIP](entrezgene:10628)
15. [DDIT3](entrezgene:1649)
16. [PHGDH](entrezgene:26227)
17. [CEBPB](entrezgene:1051)
18. [MAPK1](entrezgene:5579)
19. [FANCA](entrezgene:2177)
20. [SOCS1](entrezgene:8651)
21. [ANXA2](entrezgene:302)
22. [TCF3](entrezgene:6925)
23. [PTBP2](entrezgene:26191)
24. [PLA2G1B](entrezgene:5376)
25. [FTL](entrezgene:2495)
26. [AARS1](entrezgene:16)
27. [FANCC](entrezgene:2176)
28. [RAB1A](entrezgene:60674)
29. [BCKDK](entrezgene:8945)
30. [IGFBP2](entrezgene:3485)
31. [TPM1](entrezgene:7168)
32. [COPA](entrezgene:1314)
33. [RBM4](entrezgene:6016)
34. [KCNB1](entrezgene:4131)
35. [PTPRO](entrezgene:7494)
36. [MDH1](entrezgene:4192)
37. [SRF](entrezgene:6688)
38. [ABO](entrezgene:28)
39. [AAR2](entrezgene:22848)
40. [NCOR1](entrezgene:9463)
41. [CCNT2](entrezgene:8239)
42. [RIPK2](entrezgene:8767)
43. [SOCS5](entrezgene:8835)
44. [PDIA4](entrezgene:9361)
45. [SMG1](entrezgene:8826)
46. [SEC61A1](entrezgene:6520)
47. [DYNC1H1](entrezgene:2617)
48. [ITPR3](entrezgene:3708)
49. [PMPCA](entrezgene:29968)
50. [ATF4](entrezgene:22926)
51. [LAMTOR1](entrezgene:11151)
52. [ACAT2](entrezgene:123)
53. [LYPLA1](entrezgene:5106)
54. [DAP3](entrezgene:1601)

55. [CHERP](entrezgene:57142)
56. [HOOK3](entrezgene:30011)
57. [VIPR2](entrezgene:10797)
58. [S100A10](entrezgene:6282)
59. [STIP1](entrezgene:11168)
60. [ABCC4](entrezgene:9869)
61. [ANXA4](entrezgene:306)
62. [CAAP1](entrezgene:51441)
63. [CXCL16](entrezgene:10241)
64. [TNIP2](entrezgene:7305)
65. [HAT1](entrezgene:3021)
66. [SOX11](entrezgene:8848)
67. [CLIC1](entrezgene:1192)
68. [ARG2](entrezgene:384)
69. [LYPD3](entrezgene:11156)
70. [RET](entrezgene:5978)
71. [TEKT4](entrezgene:83667)
72. [SIRT6](entrezgene:51547)
73. [CD47](entrezgene:963)
74. [FCGRT](entrezgene:10209)
75. [HSP27](entrezgene:3151)
76. [GLUL](entrezgene:3092)
77. [SERPINB1](entrezgene:5720)
78. [SIRT5](entrezgene:51520)
79. [CNIH4](entrezgene:29117)
80. [RPL29](entrezgene:6171)
81. [ROCK2](entrezgene:11001)
82. [TBXA2R](entrezgene:2673)
83. [SNAP29](entrezgene:4637)
84. [RPRD1A](entrezgene:25932)
85. [COPS3](entrezgene:1317)
86. [ITPKB](entrezgene:3709)
87. [ZNF211](entrezgene:125144)
88. [ZCCHC7](entrezgene:9709)
89. [DNAJB5](entrezgene:843)
90. [ACTR5](entrezgene:83442)
91. [TMF1](entrezgene:7188)
92. [PRMT1](entrezgene:10109)
93. [DESI1](entrezgene:8905)
94. [RAB18](entrezgene:64428)
95. [SERPINB5](entrezgene:5723)
96. [CBY1](entrezgene:81858)
97. [ECHS1](entrezgene:170954)
98. [RFXAP](entrezgene:5934)
99. [LPAR1](entrezgene:1182)
100. [CENPH](entrezgene:64946)
101. [ACSL5](entrezgene:3140)
102. [PQLC1](entrezgene:84142)
103. [PLPPR2](entrezgene:196527)
104. [SLC39A4](entrezgene:54014)
105. [GALNT5](entrezgene:23417)
106. [IFRD1](entrezgene:3475)
107. [NUPL2](entrezgene:23127)
108. [TBC1D7](entrezgene:51669)
109. [POLR3C](entrezgene:5528)
110. [RNF145](entrezgene:55450)
111. [GID4](entrezgene:147111)
112. [ASIC2](entrezgene:80223)
113. [ECH1](entrezgene:1891)
114. [KLHL18](entrezgene:160428)
115. [FRAT2](entrezgene:6904)
116. [SLC25A39](entrezgene:84707)
117. [PRDX3](entrezgene:4848)
118. [SGPP1](entrezgene:9881)
119. [HNRNPDL](entrezgene:27069)
120. [DOLK](entrezgene:79621)
121. [ADCY9](entrezgene:118)
122. [SLC25A15](entrezgene:29091)
123. [DGKD](entrezgene:10902)
124. [ARL1](entrezgene:374383)
125. [MIS18A](entrezgene:84706)
126. [ZMPSTE24](entrezgene:4116)
127. [TBX21](entrezgene:6917)
128. [TRIM66](entrezgene:11135)
129. [FBN3](entrezgene:116028)
130. [AGPAT5](entrezgene:55207)
131. [XPO4](entrezgene:9213)
132. [TICAM2](entrezgene:54902)
133. [PTMS](entrezgene:5763)
134. [PPIL1](entrezgene:22930)
135. [IMPDH2](entrezgene:27032)
136. [NADSYN1](entrezgene:55729)
137. [AGK](entrezgene:9550)
138. [BDH1](entrezgene:122704)
139. [ZFYVE21](entrezgene:79797)
140. [MARVELD1](entrezgene:55700)
141. [SRGN](entrezgene:9061)
142. [SERPINB2](entrezgene:5711)
143. [RAB19](entrezgene:27314)
144. [ITGB3BP](entrezgene:23420)
145. [SEC11C](entrezgene:23478)
146. [CACYBP](entrezgene:51251)
147. [RGR](entrezgene:2787)
148. [MRPL37](entrezgene:55013)
149. [CCDC84](entrezgene:100129387)
150. [SNORD72](entrezgene:643911)
151. [SNORD72](entrezgene:282809)
152. [IRAK1](entrezgene:4033)
153. [LAMTOR2](entrezgene:27072)
154. [TMEM143](entrezgene:55748)
155. [VPS26B](entrezgene:55652)
156. [TMEM33](entrezgene:115677)
157. [IPKB](entrezgene:9778)
158. [RABL3](entrezgene:253980)
159. [SLC25A44](entrezgene:92255)
160. [LIPT2](entrezgene:79647)
161. [C1orf35](entrezgene:3104)
162. [ZNF410](entrezgene:10914)
163. [SMDT1](entrezgene:56731)
164. [CINP](entrezgene:280636)
165. [DDX5](entrezgene:1657)
166. [ADRM1](entrezgene:10263)
167. [RAB33B](entrezgene:85444)
168. [CWC15](entrezgene:23132)
169. [UCHL5](entrezgene:80011)
170. [OPTC](entrezgene:27236)
171. [ATAD3B](entrezgene:84904)
172. [SEPP1](entrezgene:8721)
173. [EEPD1](entrezgene:22905)
174. [MIR4712](entrezgene:101927480)
175. [FOXP1](entrezgene:22950)
176. [TRIM11](entrezgene:55631)
177. [C1orf131](entrezgene:387263)
178. [RPAP3](entrezgene:51110)
179. [DUSP1](entrezgene:5877)
180. [NDUFA13](entrezgene:11179)
181. [RSL24D1](entrezgene:150759)
182. [CCDC142](entrezgene:55266)
183. [RAB39B](entrezgene:222236)
184. [GLT8D2](entrezgene:122553)
185. [FBXO28](entrezgene:84105)
186. [SPSB4](entrezgene:153768)
187. [TSEN15](entrezgene:51657)
188. [ERGIC1](entrezgene:56954)
189. [PNKP](entrezgene:55230)
190. [RCBTB1](entrezgene:55954)
191. [TCF25](entrezgene:7769)
192. [ZNF644](entrezgene:55861)
193. [SNCAIP](entrezgene:7627)
194. [CFAP20](entrezgene:58486)
195. [DNAJC19](entrezgene:246269)
196. [DESI2](entrezgene:26071)
197. [DZIP1L](entrezgene:159013)
198. [ZNF705E](entrezgene:728855)
199. [LOC102724701](entrezgene:102724701)
200. [TMSB4XP4](ensembl:ENSG00000278384)

This list represents the top 200 targets from the focal graph search results, ordered by their relevance to the query. Each target is identified by its Entrez Gene ID, which can be used for further analysis or investigation. The targets span a wide range of biological functions and pathways, potentially offering various insights into the biological processes related to your original query.

**AI Guide Agent: Thank you for the analysis. Now, let's proceed with our research using these 200 gene symbols as the gene list to run a second focal graph search.**
Step 4: Perform a focal graph search with the identified targets
We'll use the top 200 targets identified from the [CTNNB1](https://gwps.wi.mit.edu/?chosen_gene=ENSG00000168036 "gwps:1937_CTNNB1_P1P2_ENSG00000168036|up") Perturb-seq profile to run a new focal graph search. This will help us identify potential novel targets related to the Wnt pathway and oncology.

**AI Guide Agent: Analyze the results, focusing on the profiling categories such as RNA Profiles, Proteomic Profiles, and Perturb Seq. Identify gene perturbations that show up near the top of these categories and/or are observed multiple times within a category or across categories. Consider the known targets of compounds that show up in the proteomic profile results and assess whether these compounds share a common target, especially for those compounds at the top of the list. Look for targets that appear near the top and/or are found multiple times either within or across the profiling categories. Also, assess whether the top targets might be related, e.g., are members of the same protein complex or pathway. Nominate candidate target(s) that appear to have the strongest evidence, providing the supporting data for each nomination. Evaluate their novelty, highlighting genes that are not commonly associated with the Wnt pathway. These represent putative novel gene targets for the Wnt pathway and should be considered for further investigation.**

**AI Analyst Agent: Certainly! I'll analyze the results of our focal graph search, focusing on the profiling categories as you've requested. To do this, I'll need to fetch the results from our previous search. Let's start by retrieving the data from the relevant categories. Now let's analyze them to identify potential novel targets for the Wnt pathway.**
1. RNA Profiles (GDS category):
The top results in the RNA Profiles category show several interesting gene perturbations:
a) [SSB](gpsa:D10640_SSB_UP) (up-regulated)
b) [PPP1R10](gpsa:D10549_PPP1R10_DOWN) (down-regulated)
c) [KHSRP](gpsa:D21282_KHSRP_DOWN) (down-regulated)
d) [KPNB1](gpsa:D10487_KPNB1_DOWN) (down-regulated)
e) [POLR2A](gpsa:D10545_POLR2A_DOWN) (down-regulated)

2. Perturb Seq (GWPS category):
The top results in the Perturb Seq category reveal several interesting gene perturbations:
a) [CTNNB1](https://gwps.wi.mit.edu/?chosen_gene=ENSG00000168036 "gwps:1937_CTNNB1_P1P2_ENSG00000168036|up") (up-regulated)
b) [OXA1L](gwps:6006_OXA1L_P1P2_ENSG00000155463|up) (up-regulated)
c) [EIF2S1](gwps:2541_EIF2S1_P1P2_ENSG00000134001|up) (up-regulated)
d) [EIF2B2](gwps:2536_EIF2B2_P1P2_ENSG00000119718|up) (up-regulated)
e) [EIF2B3](gwps:2537_EIF2B3_P1P2_ENSG00000070785|up) (up-regulated)

Analysis and Candidate Target Nominations:
**1. CTNNB1**: This gene is a well-known component of the Wnt signaling pathway. Its presence at the top of the Perturb Seq results validates our search strategy. However, it's not a novel target for the Wnt pathway.
**2. EIF2 Complex Members:** Several components of the eIF2 (eukaryotic initiation factor 2) complex appear in the top Perturb Seq results: [EIF2S1], [EIF2B2], [EIF2B3], [EIF2B4], and [EIF2B5]. This suggests a potential link between protein synthesis regulation and Wnt signaling.
Nomination: The eIF2 complex as a potential novel target for modulating Wnt signaling.

Supporting evidence: Multiple components of the complex appear in the top Perturb Seq results, suggesting a consistent effect.
Novelty: While protein synthesis is known to be important for cell signaling, the specific role of eIF2 in Wnt signaling is not well-established, making this a potentially novel connection.
**3. OXA1L**: This gene encodes a mitochondrial inner membrane protein involved in the insertion of mitochondrial and nuclear-encoded proteins into the mitochondrial inner membrane.
Nomination: OXA1L as a potential novel target for modulating Wnt signaling.
Supporting evidence: It appears high in the Perturb Seq results, suggesting a strong connection to the Wnt pathway perturbation profile.
Novelty: The connection between mitochondrial protein insertion and Wnt signaling is not well-established, making this a potentially novel target.
**4. SSB (Sjogren syndrome antigen B):**
This gene encodes a protein involved in RNA processing and is associated with autoimmune disorders.
Nomination: SSB as a potential novel target for modulating Wnt signaling.
Supporting evidence: It appears at the top of the RNA Profiles results, indicating a strong transcriptional response related to Wnt pathway perturbation.
Novelty: The connection between SSB and Wnt signaling is not well-established, making this a potentially novel target.
**5. PPP1R10 (Protein Phosphatase 1 Regulatory Subunit 10):**
This gene is involved in regulation of chromatin structure and gene expression.
Nomination: PPP1R10 as a potential novel target for modulating Wnt signaling.
Supporting evidence: It appears high in the RNA Profiles results, suggesting a strong transcriptional response related to Wnt pathway perturbation.
Novelty: While protein phosphatases are known to be involved in various signaling pathways, the specific role of PPP1R10 in Wnt signaling is not well-established, making this a potentially novel connection.
In conclusion, our analysis has identified several potential novel targets for modulating the Wnt pathway. The eIF2 complex members, OXA1L, SSB, and PPP1R10 stand out as the most promising candidates due to their strong representation in the profiling results and their potential novelty in relation to Wnt signaling. Further investigation into these targets could provide new insights into Wnt pathway regulation and potentially lead to new therapeutic approaches for diseases involving Wnt signaling dysregulation.

**AI Guide Agent: Thank you for this comprehensive analysis. Based on these results, we have identified several promising novel targets for modulating the Wnt pathway in oncology. Let's proceed with further investigation of our top candidates.**
<u>Step 5: Further characterization of top candidates</u>
We'll focus on our top three novel candidates:
**1. The eIF2 complex (specifically [EIF2S1], [EIF2B2], and [EIF2B3])**
**2. OXA1L**
**3. SSB**
For each of these candidates, we'll perform additional focal graph searches to gather more information and supporting evidence.
Let's start with the eIF2 complex. We'll run a focal graph search using [EIF2S1], [EIF2B2], and [EIF2B3] as our input.

**AI Guide Agent: Analyze the results, focusing on connections to Wnt signaling, cancer, and potential mechanisms of action. Look for supporting evidence in categories like Publications, Patents, and Pathways. Identify any known drugs or compounds that target these genes. Summarize the findings, highlighting the most relevant information for our investigation into novel oncology targets in the Wnt pathway.**
**AI Analyst Agent: Certainly! I'll analyze the results focusing on connections to Wnt signaling, cancer, and potential mechanisms of action. I'll look for supporting evidence in categories like Publications, Patents, and Pathways, and identify any known drugs or compounds targeting these genes. Then, I'll summarize the findings, highlighting the most relevant information for our investigation into novel oncology targets in the Wnt pathway.**
To begin, I'll need to fetch the results from our previous search. Since we don't have specific focal graph IDs mentioned in the current context, I'll start by fetching the top results from relevant categories. Let's begin with the

target category, as it's likely to contain genes related to Wnt signaling and cancer. Now, let's analyze each of these genes for their connections to Wnt signaling, cancer, and potential mechanisms of action. We'll start by fetching more detailed information about each gene.
Now that we have detailed information about these genes, let's analyze their connections to Wnt signaling, cancer, and potential mechanisms of action. We'll start by looking for publications related to each gene. Now that we have gathered information about the top targets and their related publications, let's analyze the findings focusing on connections to Wnt signaling, cancer, and potential mechanisms of action.

1. [EIF2S1](eukaryotic translation initiation factor 2 subunit alpha):
  - No direct connections to Wnt signaling were found in the top publications.
  - Cancer relevance:
    - [EIF2S1] is involved in the integrated stress response, which is important in cancer cell survival and adaptation to stress conditions.
    - Supporting evidence shows that [EIF2S1] phosphorylation is a key event in stress response pathways, which can be relevant in cancer progression and therapy resistance.
  - Potential mechanism of action:
    - [EIF2S1] phosphorylation inhibits protein synthesis by reducing the activity of eIF2B, a guanine nucleotide exchange factor.
    - The protein HRD1 has been shown to protect renal tubular epithelial cells from apoptosis by mediating [EIF2S1] ubiquitylation and degradation, suggesting a potential therapeutic target for renal diseases.

2. [EIF2B2](eukaryotic translation initiation factor 2B subunit beta):
  - No direct connections to Wnt signaling were found in the top publications.
  - Cancer relevance:
    - [EIF2B2] is part of the eIF2B complex, which is crucial for protein synthesis regulation and cellular stress response, both of which are important in cancer biology.
  - Potential mechanism of action:
    - [EIF2B2] is part of the regulatory subcomplex of eIF2B that mediates inhibition by phosphorylated eIF2α.
    - The eIF2B complex acts as a GDP dissociation stimulator protein, playing a key role in translation initiation.

3. [EIF2B3] (eukaryotic translation initiation factor 2B subunit gamma):
  - No direct connections to Wnt signaling were found in the top publications.
  - Cancer relevance:
    - Like [EIF2B2], [EIF2B3] is part of the eIF2B complex, which is important in protein synthesis regulation and stress response in cancer cells.
  - Potential mechanism of action:
    - [EIF2B3] is part of the eIF2B complex, which acts as a GDP dissociation stimulator protein.
    - The eIF2B complex plays a crucial role in the integrated stress response, which is relevant to cancer cell survival and adaptation.

While the direct connections to Wnt signaling are not apparent in the top publications for these genes, their involvement in protein synthesis regulation and stress response pathways suggests potential indirect connections to cancer progression and therapy resistance. To further investigate potential links to Wnt signaling and identify known drugs or compounds targeting these genes, let's look for pathway information and compound associations.

Based on our analysis of the top targets, related publications, pathways, and compounds, here's a summary of our findings for novel oncology targets in the Wnt pathway:

1. [EIF2S1] (eIF2α):
  - While not directly linked to Wnt signaling, [EIF2S1] plays a crucial role in the integrated stress response, which is important in cancer cell survival and adaptation.
  - [EIF2S1] phosphorylation is a key event in stress response pathways, potentially relevant to cancer progression and therapy resistance.

- Several compounds (unichem:187710733), (unichem:187721667), (unichem:187721668), (unichem:187721666), (unichem:187710671)) have been associated with [EIF2S1], suggesting potential for drug development.

2. [EIF2B2] (eIF2B subunit beta):
   - Part of the eIF2B complex, crucial for protein synthesis regulation and cellular stress response.
   - No direct compounds were identified targeting [EIF2B2], but its role in the eIF2B complex suggests it could be a potential target for modulating stress response in cancer cells.
3. [EIF2B3](eIF2B subunit gamma):
   - Also part of the eIF2B complex, important in protein synthesis regulation and stress response in cancer cells.
   - One compound, [Pcbp2-gpx4](unichem:177335995), was associated with [EIF2B3], suggesting potential for targeted drug development.

While these targets are not directly linked to Wnt signaling based on the available information, they play crucial roles in protein synthesis regulation and stress response pathways, which are important in cancer biology. Their involvement in these processes suggests potential indirect connections to cancer progression and therapy resistance, which could be relevant to Wnt pathway-related oncology research.

To further investigate novel oncology targets in the Wnt pathway, we should consider the following next steps:
1. Expand the search to include genes known to be involved in Wnt signaling and analyze their interactions with the translation initiation factors identified here.
2. Investigate the potential crosstalk between stress response pathways (involving EIF2S1, EIF2B2, and EIF2B3) and Wnt signaling in cancer cells.
3. Explore the effects of modulating these translation initiation factors on Wnt pathway activity in cancer cell lines or animal models.
4. Analyze gene expression data from cancer samples to identify correlations between the expression of these translation initiation factors and known Wnt pathway components.
5. Investigate the compounds associated with [EIF2S1] and [EIF2B3] for their potential effects on Wnt signaling and cancer cell survival.
6. Consider developing combination therapies that target both the stress response pathway (through these translation initiation factors) and the Wnt pathway to potentially enhance cancer treatment efficacy.

In conclusion, while our initial analysis didn't reveal direct connections between these translation initiation factors and Wnt signaling, their importance in cancer cell survival and stress response mechanisms suggests they could be valuable targets for novel oncology therapies. Further research is needed to elucidate potential connections between these pathways and to develop targeted approaches for cancer treatment.

**AI Guide Agent: Thank you for this detailed analysis. It's clear that while the eIF2 complex members don't have a direct known connection to Wnt signaling, they play crucial roles in processes that are relevant to cancer biology. Let's continue our investigation with our next candidate, [OXA1L](https://www.ncbi.nlm.nih.gov/gene/5018 "entrezgene:5018").**

**AI Guide Agent: Analyze the results, focusing on connections to Wnt signaling, cancer, and potential mechanisms of action. Look for supporting evidence in categories like Publications, Patents, and Pathways. Identify any known drugs or compounds that target this gene. Summarize the findings, highlighting the most relevant information for our investigation into novel oncology targets in the Wnt pathway.**

**AI Analyst Agent: Certainly! I'll analyze the results for connections to Wnt signaling, cancer, and potential mechanisms of action, focusing on supporting evidence from various categories. I'll also look for known drugs or compounds targeting this gene. To do this effectively, I'll need to fetch more detailed information about the gene and its relationships. Let's start by retrieving the entity information and exploring its connections.**

**AI Analyst Agent: Thank you for providing the entity information for [OXA1L](https://www.ncbi.nlm.nih.gov/gene/5018 "entrezgene:5018"). Now, let's explore its connections and gather more information about its relationships to Wnt signaling, cancer, and potential mechanisms of action. Now, Let's analyze the results, focusing on connections to Wnt signaling, cancer, and potential mechanisms of action. We'll also look for any known drugs or compounds that target this gene.**

1. Connections to Wnt signaling:
Unfortunately, none of the retrieved publications directly mention Wnt signaling in relation to [OXA1L]. This suggests that there might not be a strong direct connection between [OXA1L] and Wnt signaling based on the current literature.

2. Connections to cancer:
While [OXA1L] is not directly implicated in cancer in most of these publications, there are some relevant findings:
a. [Genome-wide CRISPR screen identifies](pubmed:30093494) mentions a genome-wide CRISPR screen, which could potentially identify genes involved in various cellular processes, including cancer-related pathways.
b. [Genome-wide CRISPR screening reveals nucleotide synthesis negatively regulates autophagy](pubmed:34000301) discusses autophagy regulation, which is known to play a role in cancer development and progression.
c. [The Mitochondrial Transacylase, Tafazzin, Regulates for AML Stemness by Modulating Intracellular Levels of Phospholipids](pubmed:30930145) focuses on acute myeloid leukemia (AML) and the role of mitochondrial proteins in cancer stem cells. Although this study doesn't directly involve [OXA1L], it demonstrates the importance of mitochondrial proteins in cancer biology.
d. [Mitochondrial carrier homolog 2 is necessary for AML survival](pubmed:32299104) also discusses the role of mitochondrial proteins in AML, further highlighting the potential importance of mitochondrial function in cancer.

3. Potential mechanisms of action:
[OXA1L] appears to be primarily involved in mitochondrial function and protein insertion into the inner mitochondrial membrane. Key findings include:
a. [Topogenesis of mammalian Oxa1, a component of the mitochondrial inner membrane protein export machinery](pubmed:19349278) describes [OXA1L] as a crucial component of the mitochondrial inner membrane protein export machinery.
b. [Properties of the C-terminal tail of human mitochondrial inner membrane protein Oxa1L and its interactions with mammalian mitochondrial ribosomes](pubmed:20601428) reveals that [OXA1L] interacts with mitochondrial ribosomes, suggesting a role in co-translational insertion of mitochondria-synthesized proteins.
c. [Identification of TMEM126A as OXA1L-interacting protein reveals cotranslational quality control in mitochondria](pubmed:38199007) demonstrates that [OXA1L] is involved in protein insertion and quality control within mitochondria.
These mechanisms suggest that [OXA1L] plays a crucial role in maintaining mitochondrial function and protein homeostasis, which could indirectly affect various cellular processes, including those related to cancer.

4. Known drugs or compounds targeting [OXA1L]:
Based on the provided publications, there are no specific drugs or compounds mentioned that directly target [OXA1L]. However, the importance of mitochondrial function in cancer, as highlighted in some of the studies, suggests that targeting mitochondrial proteins like [OXA1L] could potentially be a novel approach for cancer treatment.

Summary and relevance to novel oncology targets in the Wnt pathway:
1. [OXA1L] is a crucial component of the mitochondrial inner membrane protein export machinery and plays a role in protein insertion and quality control within mitochondria.
2. While there is no direct evidence linking [OXA1L] to Wnt signaling, its importance in mitochondrial function could potentially have indirect effects on cellular processes regulated by Wnt signaling.
3. The role of mitochondrial proteins in cancer, particularly in AML, is highlighted in several studies. This suggests that targeting mitochondrial function could be a promising approach for cancer treatment.

4. [OXA1L] itself is not directly implicated as an oncology target based on the current literature. However, its involvement in crucial mitochondrial processes makes it an interesting candidate for further investigation in the context of cancer biology.
5. The lack of known drugs or compounds targeting [OXA1L] suggests that it could be a novel target for drug development if its relevance to cancer biology is established through further research.

To further explore [OXA1L] as a potential novel oncology target in the Wnt pathway, additional research would be needed to:
1. Investigate possible interactions between [OXA1L] and components of the Wnt signaling pathway.
2. Examine the expression levels and potential mutations of [OXA1L] in various cancer types, particularly those associated with aberrant Wnt signaling.
3. Conduct functional studies to determine if modulation of [OXA1L] activity affects Wnt signaling or cancer cell behavior.
4. Explore the potential indirect effects of [OXA1L] on Wnt signaling through its role in mitochondrial function and cellular metabolism.

While [OXA1L] may not be directly linked to Wnt signaling based on the current literature, its crucial role in mitochondrial function and the emerging importance of mitochondrial proteins in cancer biology make it an interesting candidate for further investigation in the context of novel oncology targets.

**AI Guide Agent: Thank you for this thorough analysis of [OXA1L]. While it doesn't show a direct connection to Wnt signaling, its role in mitochondrial function and potential indirect effects on cancer biology make it an interesting candidate for further investigation. Let's move on to our final candidate, [SSB](https://www.ncbi.nlm.nih.gov/gene/6741 "entrezgene:6741").**

**AI Guide Agent: Analyze the results, focusing on connections to Wnt signaling, cancer, and potential mechanisms of action. Look for supporting evidence in categories like Publications, Patents, and Pathways. Identify any known drugs or compounds that target this gene. Summarize the findings, highlighting the most relevant information for our investigation into novel oncology targets in the Wnt pathway.**

**AI Analyst Agent: Certainly! To analyze the results and gather the information you've requested, I'll need to perform a series of focal graph searches and data retrievals. Let's start by fetching the search results and then we'll analyze the connections to Wnt signaling, cancer, and potential mechanisms of action. It looks like we have one primary target to focus on: [SRF](https://www.ncbi.nlm.nih.gov/gene/6741 "entrezgene:6741"). Let's gather more information about this gene and its connections to Wnt signaling, cancer, and potential mechanisms of action.**
**Now that we have more information about [SSB], let's analyze its connections to Wnt signaling, cancer, and potential mechanisms of action. We'll start by fetching results from relevant categories.**
**Now, let's analyze the results and summarize the findings:**

1. Gene Information:
[SSB] (Sjögren syndrome antigen B) is also known as La/SSB or La autoantigen. It is a small RNA binding exonuclease protection factor. The gene is involved in various cellular processes, including RNA polymerase III transcription termination.

2. Connection to Wnt signaling:
Based on the pathway results, there is no direct evidence of [SSB] being involved in Wnt signaling. However, this doesn't rule out the possibility of indirect connections or less well-characterized interactions.

3. Connection to cancer:
While [SSB] is not primarily known as an oncogene or tumor suppressor, there is evidence of its involvement in cancer-related processes:

a. [A study published in 2022](pubmed:35271311) investigated the role of [SSB] in hepatocellular carcinoma (HCC). The research suggests that [SSB] may play a role in HCC progression.
b. [Another study from 2022](pubmed:35831314) examined the potential of [SSB] as a biomarker for colorectal cancer.
c. [Research published in 2021](pubmed:33961781) explored the role of [SSB] in gastric cancer, suggesting it might be involved in cancer cell proliferation and metastasis.

4. Potential mechanisms of action:
Based on the available information, [SSB] appears to have several potential mechanisms of action that could be relevant to cancer:
a. RNA processing and stability: As a small RNA binding protein, [SSB] may regulate the stability and processing of various RNAs, which could affect gene expression in cancer cells.
b. Transcription regulation: [SSB] is involved in RNA polymerase III transcription termination, suggesting it may play a role in regulating gene expression at the transcriptional level.
c. Autoimmune responses: Given its role as an autoantigen in Sjögren's syndrome, [SSB] may be involved in modulating immune responses, which could be relevant in cancer immunology.

5. Known drugs or compounds:
The search results don't provide information about specific drugs or compounds targeting [SSB]. However, the presence of patents suggests that there may be ongoing efforts to develop therapeutics related to this gene:
a. [A patent from 2023](patents:48044435) appears to be the most relevant, potentially involving [SSB] in a therapeutic context.
b. [Several other patents](patents:41398504) from 2022 and 2023 also mention [SSB], suggesting ongoing research and development efforts.

Summary:
While [SSB] is not directly linked to Wnt signaling based on the provided results, it shows potential as a novel oncology target. Its involvement in RNA processing, transcription regulation, and potential roles in various cancers (including hepatocellular carcinoma, colorectal cancer, and gastric cancer) make it an interesting candidate for further investigation. The presence of recent publications and patents suggests ongoing research interest in [SSB] as a potential therapeutic target.

To further explore [SSB] as a novel oncology target, consider the following steps:
1. Investigate its potential indirect interactions with Wnt signaling components.
2. Explore its role in regulating cancer-related genes and pathways.
3. Examine its expression levels and mutations in various cancer types.
4. Investigate its potential as a biomarker or therapeutic target in different cancers.
5. Look into ongoing clinical trials or preclinical studies targeting [SSB] or its related pathways.

These findings suggest that [SSB] could be a promising area for further research in oncology, even if its connection to Wnt signaling is not immediately apparent from the current results.

**AI Guide Agent: Thank you for this comprehensive analysis of our three candidate targets: the eIF2 complex ([EIF2S1], [EIF2B2], and [EIF2B3]), [OXA1L], and [SSB]. Let's summarize our findings and outline potential next steps for our research program.**

Summary of Findings:

**1. eIF2 Complex:**
- Involved in protein synthesis regulation and cellular stress response.
- Potential indirect connections to cancer progression and therapy resistance.
- Several compounds associated with [EIF2S1], suggesting drug development potential.
- No direct link to Wnt signaling found, but potential for crosstalk between stress response and Wnt pathways.

**2. OXA1L:**
- Crucial component of mitochondrial inner membrane protein export machinery.
- No direct connection to Wnt signaling found.
- Potential indirect effects on cancer biology through mitochondrial function.
- No known drugs targeting [OXA1L], suggesting novelty as a potential target.

**3. SSB:**
- Involved in RNA processing, transcription regulation, and autoimmune responses.
- No direct connection to Wnt signaling found.
- Evidence of involvement in hepatocellular carcinoma, colorectal cancer, and gastric cancer.
- Recent patents suggest ongoing research interest as a potential therapeutic target.

Next Steps:

1. Further investigation of eIF2 complex:
   - Explore potential crosstalk between stress response pathways and Wnt signaling in cancer cells.
   - Analyze gene expression data to identify correlations between eIF2 complex components and Wnt pathway genes in cancer samples.
   - Investigate the effects of modulating eIF2 complex components on Wnt pathway activity in cancer cell lines or animal models.

2. OXA1L research:
   - Examine expression levels and potential mutations of [OXA1L] in cancers associated with aberrant Wnt signaling.
   - Conduct functional studies to determine if modulation of [OXA1L] activity affects Wnt signaling or cancer cell behavior.
   - Explore potential indirect effects of [OXA1L] on Wnt signaling through its role in mitochondrial function and cellular metabolism.

3. SSB exploration:
   - Investigate potential indirect interactions between [SSB] and Wnt signaling components.
   - Explore its role in regulating cancer-related genes and pathways, particularly in hepatocellular carcinoma, colorectal cancer, and gastric cancer.
   - Examine its potential as a biomarker or therapeutic target in different cancer types.

4. Integrative analysis:
   - Perform pathway analysis to identify potential connections between our candidate targets and Wnt signaling components.
   - Conduct protein-protein interaction studies to uncover any direct or indirect links between our candidates and known Wnt pathway proteins.

5. Experimental validation:
   - Design in vitro experiments to test the effects of modulating our candidate targets on Wnt signaling activity and cancer cell behavior.
   - Consider developing cell line or animal models with altered expression of our candidate genes to study their impact on cancer progression and Wnt signaling.

6. Drug discovery efforts:
   - For the eIF2 complex, further investigate the identified compounds for their potential effects on Wnt signaling and cancer cell survival.
   - For [OXA1L] and [SSB], initiate small molecule screening efforts to identify potential inhibitors or modulators.

7. Collaboration and expertise expansion:
   - Consider collaborating with experts in Wnt signaling, cancer biology, and drug discovery to broaden the scope and depth of our research.
   - Explore partnerships with pharmaceutical companies or academic institutions with experience in developing therapies targeting similar pathways.

By pursuing these next steps, we can further evaluate the potential of our candidate targets as novel oncology targets in the Wnt pathway. While none of them showed direct connections to Wnt signaling in our initial analysis, their roles in important cellular processes and potential indirect effects on cancer biology make them promising candidates for further investigation.
